# Restoring the Multiple Sclerosis Associated Imbalance of Gut Indole Metabolites Promotes Remyelination and Suppresses Neuroinflammation

**DOI:** 10.1101/2024.10.27.620437

**Authors:** Larissa Jank, Saumitra S. Singh, Judy Lee, Asmita Dhukhwa, Fatemeh Siavoshi, Deepika Joshi, Veronica Minney, Kanak Gupta, Sudeep Ghimire, Pragney Deme, Vinicius A. Schoeps, Karthik Soman, Dimitrios Ladakis, Matthew Smith, Kamil Borkowski, John Newman, Sergio E. Baranzini, Emmanuelle L. Waubant, Kathryn C. Fitzgerald, Ashutosh Mangalam, Norman Haughey, Michael Kornberg, Xitiz Chamling, Peter A. Calabresi, Pavan Bhargava

## Abstract

In multiple sclerosis (MS) the circulating metabolome is dysregulated, with indole lactate (ILA) being one of the most significantly reduced metabolites. We demonstrate that oral supplementation of ILA impacts key MS disease processes in two preclinical models. ILA reduces neuroinflammation by dampening immune cell activation as well as infiltration; and promotes remyelination and *in vitro* oligodendrocyte differentiation through the aryl hydrocarbon receptor (AhR). Supplementation of ILA, a reductive indole metabolite, restores the gut microbiome’s oxidative/reductive metabolic balance by lowering circulating indole acetate (IAA), an oxidative indole metabolite, that blocks remyelination and oligodendrocyte maturation. The ILA-induced reduction in circulating IAA is linked to changes in IAA-producing gut microbiota taxa and pathways that are also dysregulated in MS. Notably, a lower ILA:IAA ratio correlates with worse MS outcomes. Overall, these findings identify ILA as a potential anti-inflammatory remyelinating agent and provide insights into the role of gut dysbiosis-related metabolic alterations in MS progression.

**One Sentence Summary:** Indole lactate, a postbiotic metabolite reduced in MS, corrects gut microbiome metabolic imbalances associated with remyelination and neuroinflammation.

## INTRODUCTION

Multiple sclerosis (MS) is a chronic demyelinating disease of the central nervous system (CNS). It is the most common non-traumatic disabling neurological disorder among young adults (*1*), yet the cause of MS is not fully understood. Environmental factors such as diet and the gut microbiome have been identified as factors contributing to the cause and progression of MS. The gut microbiome, its metabolic pathways, and circulating gut-derived metabolites are dysregulated in people with MS (pwMS) (*2*). The aromatic amino acid (AAA) metabolite indole lactate (ILA) is among the top dysregulated metabolites in pwMS (*3, 4*). Circulating ILA is lower in adult and pediatric pwMS compared to healthy controls and lower in progressive compared to age-matched relapsing-remitting MS (*5*). We leveraged these clinical observations to identify a novel and potentially safe therapeutic strategy for MS. We investigated if restoring the gut microbial metabolic imbalance through the supplementation of ILA, which is naturally present in fermented food and is exclusively produced by the gut microbiome (*6*), could dampen neuroinflammation and promote remyelination. Remyelination agents are of particular interest since FDA-approved treatments available for MS primarily target inflammatory aspects of the disease. Promoting remyelination could help restore function, especially in progressive disease during which the capacity of the CNS to spontaneously remyelinate is severely limited. Another potential benefit of ILA as a treatment is that it could be administered in a personalized medicine approach to patients with low circulating ILA levels.

In previous studies, ILA has been shown to exert anti-inflammatory effects in the periphery by reducing T-cell proliferation and pro-inflammatory cytokine release from macrophages while upregulating anti-inflammatory cytokines. ILA crosses the blood-brain barrier and is detectable in both the mouse (*6*) and human (*7*) CNS. ILA-producing bacteria reduce experimental autoimmune encephalomyelitis (EAE) severity by modulating the adaptive immune response (*8*). Interestingly, other gut-derived AAA metabolites have been shown to regulate oligodendrocyte precursor cell (OPC) differentiation (*9, 10*). Based on these findings, we assessed the effect of oral ILA supplementation on two key MS processes, neuroinflammation and remyelination, using two animal models – the EAE model and the cuprizone (CPZ) toxic demyelination model. *In vitro,* we confirmed and expanded previous findings describing ILA’s effects on immune cells and also found that ILA promotes OPC differentiation through aryl hydrocarbon receptor (AhR). Investigation of ILA’s mechanism of action revealed that ILA restores gut microbial taxa and functional pathways that are dysregulated in pwMS. This study not only provides evidence for ILA as a potential remyelinating agent with additional anti-inflammatory effects but also provides new insights into how metabolic changes in pwMS affect oligodendrocyte (OL) function and repair.

## RESULTS

### ILA supplementation attenuates EAE severity and neuroinflammation

Oral ILA supplementation (2mM ad libitum in drinking water which equates to approximately 1.2mg/day/mouse) beginning one week after EAE induction reduced both peak and endpoint clinical scores compared to vehicle (acidified water with a pH matched to ILA) (Fig. 1A). Equimolar concentrations of ILA administered parenterally had similar effects (Fig. S1A). To better represent a possible clinical treatment scenario, we focused on oral supplementation for all future experiments.

**Fig. 1.**
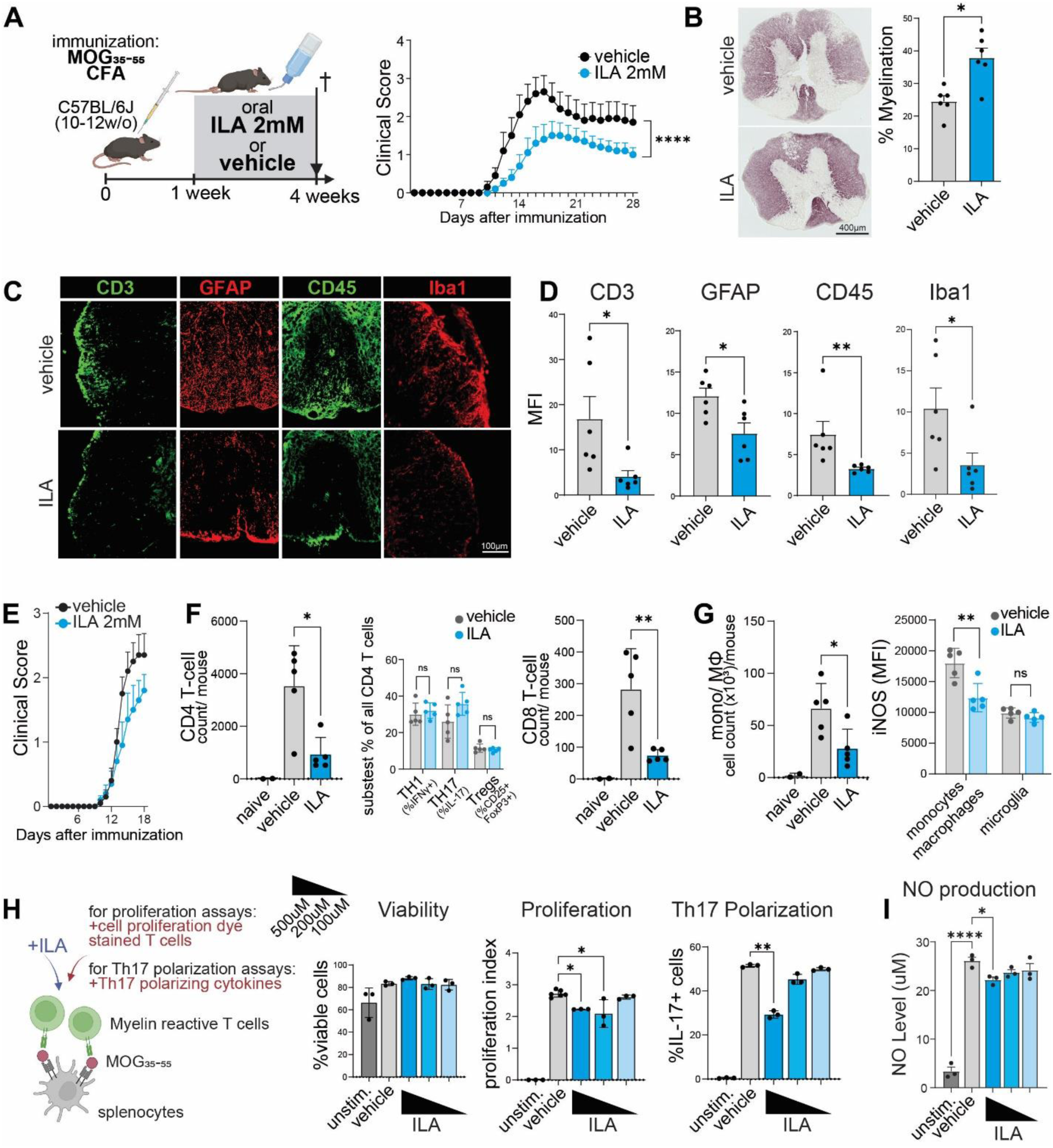
ILA supplementation attenuates EAE severity and neuroinflammation. (A) Schematic of the experimental setup (left) and Mean Hooke clinical EAE severity score of ILA- and vehicle-treated mice ±SEM (right). (B) Histopathology analysis of spinal cords with Black gold II staining after three weeks of treatment with representative images (left) and quantification of the mean % myelinated area (right). (C) Representative images of immunofluorescence staining for T-cells (CD3), leukocytes (CD45), and astrogliosis (GFAP) in the spinal cord after three weeks of treatment and (D) quantification of the mean fluorescent intensities (MFI). (E) Mean clinical scores of mice used to assess ILA’s effects on the immune infiltrate at the peak of EAE ±SEM (same setup as in A but mice were sacrificed 18 days post-immunization). (F) Flow cytometric analysis of the mean number of infiltrating CD4+ (left) and CD8+ (right) T-cells per mouse in the spinal cord at day 18 and analysis of the mean % of T-cell subsets of all infiltrating CD4+ T-cells (center). (G) Mean number (x10^3^) of infiltrating monocytes/ macrophages per mouse (left) and mean MFI of intracellular iNOS staining as a marker of cell activation (right). (H) Schematic of T-cell proliferation and Th17 polarization assays with myelin-reactive CD4+ T-cells isolated 2D2 mice and flow cytometric analysis of the mean % viability (left), mean proliferation index (center), and the mean % of IL-17+ cells i.e. Th17 polarization (right) of these T-cells treated with vehicle or 100uM-500uM ILA for the duration of these assays (3 days). (I) Griess assay measuring nitrite levels in the supernatant of peritoneal macrophages stimulated with LPS+IFNγ and treated with vehicle or 100uM-500uM ILA. The n=10 mice per group for a and 6 mice per group for C-D, 5 mice per ILA/ vehicle group for E-G, and H-I show 3 technical replicates from one of at least two independent experiments. Error bars represent ±SD, unless mentioned otherwise. Statistical significance was determined by repeated measures two-way ANOVA for EAE scoring, Mann–Whitney U test for B, D, F (center) and G (right), and one-way ANOVA with Tukey’s post-hoc test for F (remaining), G (left), H and I. (*P <0.05; **P <0.01; ****P <0.0001).

After three weeks of treatment, ILA-supplemented EAE mice had less severe spinal cord demyelination than vehicle-treated mice (Fig. 1B). At this time point, ILA treatment also reduced CD3+ and CD45+ staining, demonstrating reduced immune cell infiltration and reduced reactive astrogliosis and microgliosis/infiltrating macrophages, based on reduced GFAP and Iba-1 staining (Fig. 1C-D). At 2.5 weeks post-immunization – the peak of EAE and a time point at which ILA supplementation ameliorated clinical scores (Fig. 1E) – ILA not only decreased the number of infiltrating T-cells (Fig. 1F; ILA had no impacting relative proportion of infiltrating T-cell subsets) but also of monocytes/macrophages (Fig. 1G) in the spinal cord (gating strategy Fig. S1B-D). ILA also reduced intracellular iNOS expression in infiltrating monocytes/macrophages, suggesting reduced inflammatory activation (Fig. 1G).

*In vitro,* ILA had direct effects on T-cells and myeloid cells. ILA inhibited MOG-induced proliferation and Th17 polarization of myelin reactive T-cells in a dose-dependent manner (Fig.1H, experiments with WT T-cells Fig. S1D-E). Corroborating *in vivo* findings, ILA reduced nitric oxide (NO) production in activated primary peritoneal macrophages also in a dose-dependent manner (Fig. 1I).

### ILA enhances remyelination after cuprizone-induced demyelination

In the cuprizone (CPZ) -induced demyelination model, we tested ILA’s effects on remyelination. Mice were fed a 0.2% CPZ diet for five weeks to induce demyelination and then returned to regular feed. During the remyelination phase, mice were supplemented with either ILA or vehicle orally for two weeks (Fig. 2A). After two weeks of treatment, ILA-treated mice had larger areas of remyelination in the corpus callosum (CC) (Fig. 2B). In electron microscopy morphometric analysis of the CC, the g-ratio was reduced (denoting an increase in myelin thickness) and the percentage of myelinated axons was increased with ILA treatment compared to vehicle (Fig. 2C). In the CPZ model, reactive gliosis impacts remyelination. Using immunofluorescence staining for markers of astrogliosis (GFAP), activated microglia (Iba1 and CD68 [lysosomal marker]), and homeostatic microglia (TMEM119), we analyzed the extent of inflammation in the CC. ILA supplementation reduced GFAP, Iba1, and CD68 staining and increased TMEM119 staining, altogether suggesting a reduction in inflammatory glial activation and a return to a more homeostatic state. Further, ILA supplementation increased the percentage of mature OLs (CC1+Olig2+/Olig2+) in the CC (Fig. 2D).

**Fig. 2.**
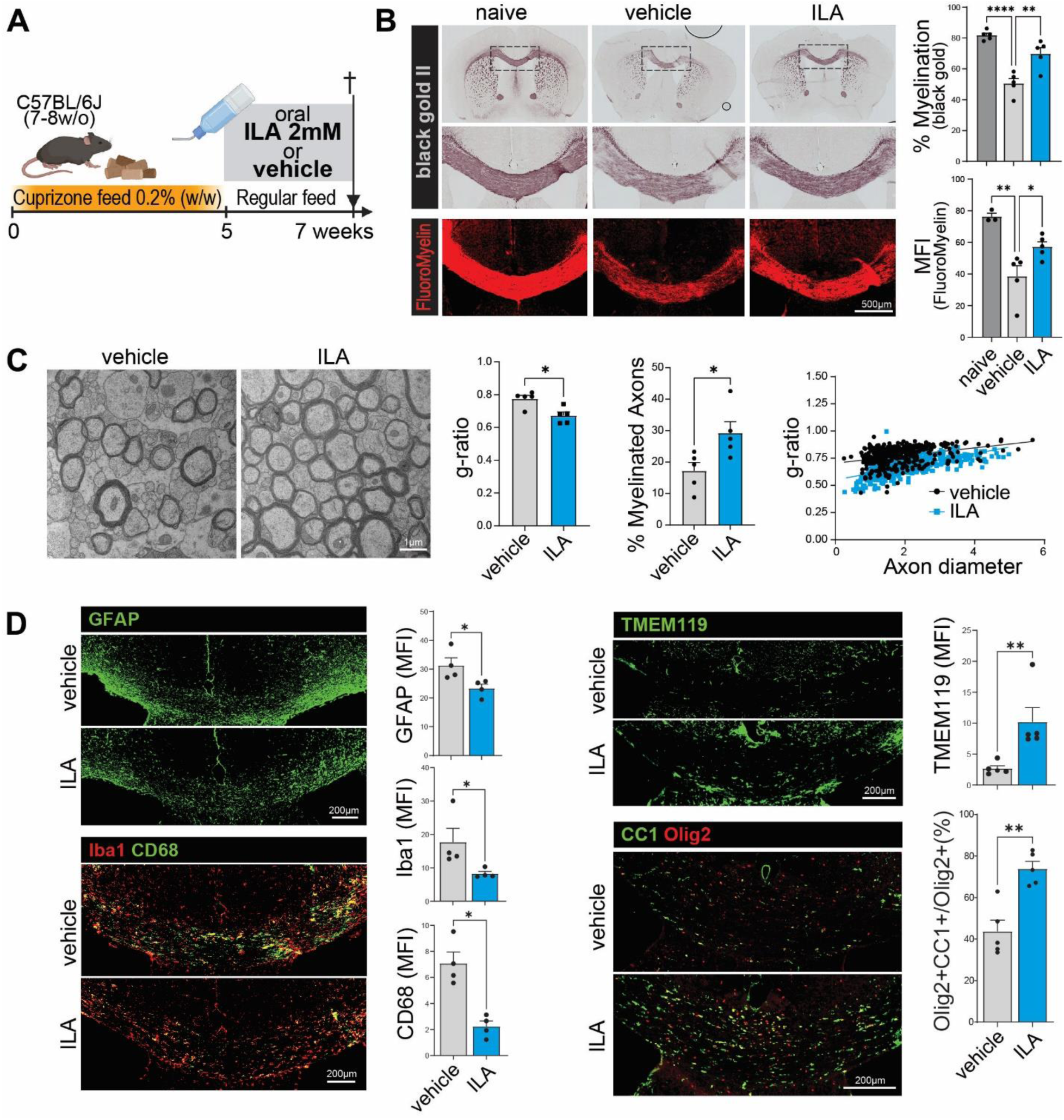
ILA promotes remyelination after cuprizone-induced demyelination. (A) Schematic of the experimental setup, where ILA/ vehicle was supplemented during the remyelination phase. (B) Histopathology analysis of the corpus callosum (CC) myelination with Black gold II and FluoroMyelin staining after two weeks of treatment with representative images (left) and quantification of the mean % myelination of the CC and MFI of the CC, respectively (right). (C) Electron microscopy morphometric analysis of the CC with representative images (left) and quantification of the mean g-ratio (center left), mean % myelinated axons (center right), and their correlation (right) obtained using the MyelTracer software. (D) Representative images of immunofluorescence staining for astrogliosis (GFAP), monocytes/macrophages and microglia (Iba1), a lysosomal marker (CD68), homeostatic microglia (TMEM119), and mature oligodendrocytes (Olig2+CC1+) in the CC after two weeks of treatment and quantification of the mean fluorescent intensities (MFI) and the % mature oligodendrocytes of all oligodendrocyte lineage cells (%Olig2+CC1+/Olig2+). The n= 5 mice per group for B-C and 4 mice per group for D. Error bars represent ±SD. Statistical significance was determined by Mann–Whitney U test for C-D, and one-way ANOVA with Dunnett’s post-hoc test for B. (*P <0.05; **P <0.01; ****P <0.0001).

### ILA promotes AhR-mediated OPC differentiation in vitro

Next, we investigated the direct effects of ILA on the differentiation of primary murine OPCs into mature OLs and observed an increased percentage of mature OLs (%CC1+Olig2+/Olig2+) in a dose-dependent manner (Fig. 3A). Using human embryonic stem cell-derived triple reporter OPC cell lines (hESC-OPCs) from two distinct donors (*11*) we confirmed that ILA also dose-dependently promotes human OPC differentiation (Fig. 3B).

**Fig. 3.**
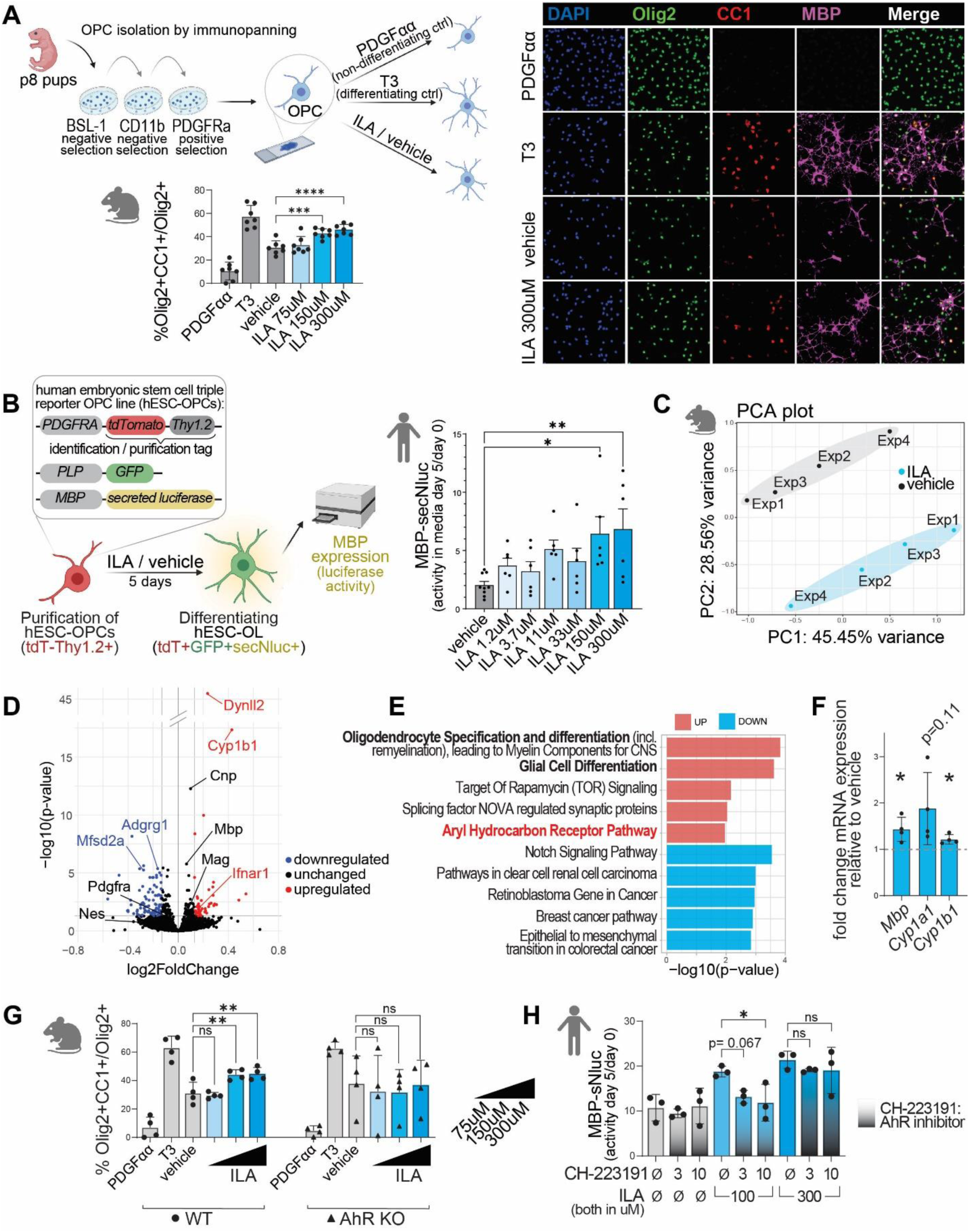
ILA promotes OPC differentiation in vitro, which is mediated by AhR. (A) Schematic of OPC differentiation assays with primary murine OPCs (top left). Immunocytochemistry of the OPCs treated with control hormones, vehicle or 75-300uM ILA for the duration of the assay with representative images of staining for a pan-oligodendrocyte marker (Olig2), oligodendrocyte maturity (CC1) and myelin proteins (MBP) (top right) and quantification of the mean % of mature oligodendrocytes of all oligodendrocytes (%Olig2+CC1+/Olig2+) (bottom left) (n= 7 independent experiments). (B) Schematic of the human embryonic stem cell triple reporter OPC cell line (hESC-OPCs) differentiation assay (left) and quantification of the change in secreted luciferase activity (measure of *MBP* transcription) in the supernatants of hESC-OPCs after 5 days of treatment with vehicle or 1.2-300uM ILA (right) (n= 6 technical replicates from one donor representative of two donors). (C) Principal component analysis on bulk RNASeq data from murine OPCs treated with vehicle or 300uM ILA (same setup as in A but collected RNA) from four independent experiments. (D) Volcano plot showing genes significantly (cutoffs: P <0.05; 5% in/decrease) upregulated (red) and downregulated (blue) with ILA treatment (genes of interest below the selected cutoffs shown in black). (E) Top 5 up- (red) and downregulated (blue) pathways determined by gene set over-representation analysis ranked by p-value (F) Confirmation of the key RNASeq findings qPCR. Shown is the mean fold-change in mRNA expression with ILA treatment compared to vehicle. (G) Quantification of the differentiation of WT (circle) and AhR-/- (triangle) primary murine OPCs into mature oligodendrocytes (%Olig2+CC1+/Olig2+) in the presence of vehicle or 75-300uM ILA (same setup as in A) (n= 4 independent experiments). (H) Quantification of the mean change in secreted luciferase activity (*MBP* transcription) in the supernatants of hESC-OPCs with vehicle or 1.2-300uM ILA treatment in the presence and absence of the competitive AhR inhibitor CH-223191 (3-10uM) (n= 3 independent experiments; cells from two donors). Error bars represent ±SD. Statistical significance was determined by one-way ANOVA with Dunnett’s post-hoc test for A, B, G, and H and by one sample t-test for F. (*P <0.05; **P <0.01).

To determine possible mechanisms through which ILA acts on OPCs, we carried out an exploratory transcriptomics experiment on primary murine OPCs treated with ILA or vehicle (same setup as for Fig. 3A). We included cells from 4 different isolations, treated on separate days, to improve the robustness of our findings, which came at the cost of slightly increased variance (Fig. 3C). Nonetheless, we observed several differentially regulated genes of interest. The AhR-responsive gene, *Cyp1b1,* was among the top upregulated genes, and other AhR-responsive genes (*Aldh3a2*, *Ifnar1)* were also upregulated. Genes expressed in differentiated OLs (*Cnp*, *Dynll2*) were upregulated while markers of OPCs (*Nes*, *Adgrg1*, *Mfsd2a*) were downregulated with ILA treatment (Fig. 3D; Supplementary Tables Sheet 1). The upregulation of OPC differentiation pathways and AhR pathways were confirmed in over-representation pathway analysis using WikiPathway gene sets (Fig. 3E; Supplementary Tables Sheet 2). The most significantly downregulated pathway was the notch pathway, which when overactivated blocks OPC differentiation (*12*). Key findings from the RNASeq experiment were confirmed with qPCR (Fig. 3F).

We tested the relevance of AhR for ILA’s effects of ILA on murine and human OPCs using murine AhR-/- OPCs and an AhR inhibitor (CH-223191). In primary murine AhR-/- OPCs, ILA failed to induce OPC differentiation (Fig. 3G). Similarly, CH-223191, a competitive AhR inhibitor (*13*), abolished ILA’s pro-differentiating effects on hESC-OPCs in a concentration-dependent manner (Fig. 3H).

### Oral ILA supplementation reduces circulating indole acetate (IAA) levels and alters the gut microbiome

In MS, gut microbial pathways are dysregulated with a shift from reductive to oxidative pathways (*3*). One reductive/oxidative AAA metabolite pair is ILA (reductive) and IAA (oxidative) (Fig. 4A). To investigate if ILA supplementation impacts this metabolic balance, we measured plasma ILA and IAA levels in ILA-supplemented mice. Interestingly, circulating IAA levels decreased with ILA supplementation both in healthy (Fig. 4A) and EAE mice (Fig. 4B-C). The largest and most significant change in EAE was in the ILA:IAA ratio. This ratio negatively correlated with the cumulative clinical EAE score (Fig. 4D).

**Fig. 4.**
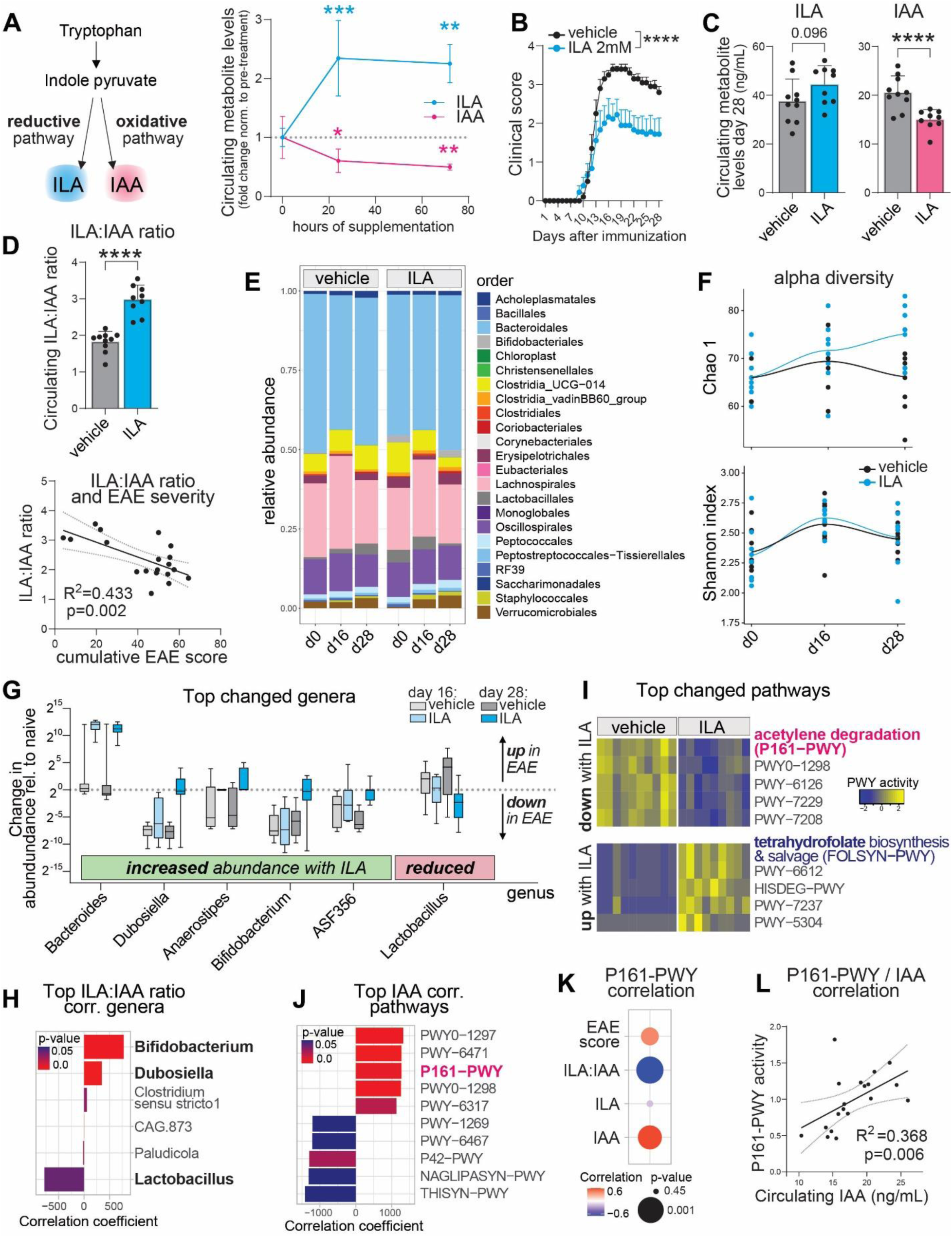
ILA supplementation reduces circulating IAA levels and alters the gut microbiome. (A) Diagram of ILA and IAA production via reductive and oxidative pathways, respectively (left) and mean fold-change ±SD in circulating ILA (blue) and IAA (pink) relative to baseline levels with oral ILA (2mM) supplementation in healthy mice (n= 5 mice per time point). (B) Mean clinical score of EAE mice ±SEM treated with ILA/ vehicle (from 1 week post-immunization until experiment completion i.e. 4 weeks) and included in metabolite and microbiome analyses in C-J (C) Mean plasma ILA and IAA levels ±SD in these mice at day 28. (D) Circulating ILA:IAA ratio (top) and its correlation with the cumulative EAE score (bottom). (E) Sequence-based bacterial analysis of fecal pellets collected before immunization, at 16 and 28 days post-immunization shows changes in beta diversity over time and with treatment. Shown are mean relative abundances of taxa (order level) for each group over time. (F) Alpha diversity analysis using Chao1 (top) and Shannon index (bottom) with the measures for each mouse plotted at each time point and a spline indicating overall change over time in the groups. (G) Top changed genera with ILA treatment compared to vehicle ranked by magnitude of change with ILA (cutoff FDR<2%). Shown is the mean change in abundance relative to baseline (pre-immunization). (H) Top ILA:IAA ratio correlated genera (cutoff P <0.05) ranked by correlation coefficient. (I) Heatmap of the top 5 most significantly up-/ down-regulated gut microbial pathways with ILA treatment compared to vehicle ranked by magnitude of change. (J) Top 5 most significantly positively and negatively IAA correlated pathways in ILA-treated mice. (K) Correlation of P161-PWY activity with circulating metabolites and the cumulative EAE score with red indicating positive and blue negative correlation. (L) Correlation of P161-PWY activity with IAA levels including both treatment groups. The n= 9-10 mice per group for B-L. Statistical significance was determined by one-way ANOVA with Dunnett’s post-hoc test for A and by Mann–Whitney U test for C and top D. (*P <0.05; **P <0.01; ***P < 0.001; ****P <0.0001). All correlations are Spearman’s rank correlations.

Since circulating IAA is primarily produced by gut bacteria (*6*), we investigated ILA’s effect on the gut microbiome. We induced EAE, then 7d post-immunization started oral ILA or vehicle supplementation in the drinking water (same set up as in Fig. 1A), and collected fecal pellets before induction (d0), at peak (d16), and at a chronic (d28) time point. In line with findings of previous studies (*14*) the alpha diversity (single sample diversity), assessed using *Chao1* and the Shannon index, mildly increased during EAE. ILA treatment had no effect on the Shannon index but increased *Chao1* compared to vehicle (*Chao1* over time for ILA R=0.56, p-value=0.003 and for vehicle R=0.02, p-value=0.94; Fig. 4F). The beta diversity (diversity across samples) also changed both during EAE and with ILA treatment (Fig. 4E, Fig. S2A). Since the beta diversity of vehicle and ILA-treated groups differed slightly at baseline (Fig. S2B), we compared changes in bacterial abundances relative to the day 0 composition. Additionally, we confirmed key analyses with an alternative approach directly comparing endpoint differences (Fig. S2D-E). ILA treatment for 4 weeks increased the abundance of *Bacteroides*, *Dubosiella, Anaerostipes*, and *Bifidobacterium* and decreased the abundance of *Lactobacillus* (Fig. 4G, Fig. S2D; Supplementary Tables Sheet 3). At the species level, ILA increased *D. newyorkensis*, *B. thetaiotaomicron* and *B. pseudolongum* while *F. rodentium* decreased (Fig. S2C). *Dubosiella* and *Bifidobacterium* were positively, and *Lactobacillus* was negatively correlated with the circulating ILA:IAA ratio (Fig. 4H).

ILA supplementation also altered the metabolic function of the gut microbiome (Fig. 4I, Fig. S2E; Supplementary Tables Sheet 4). The acetylene degradation pathway (P161-PWY), which includes reactions leading to IAA production downstream, was the most significantly downregulated pathway in ILA-treated compared to vehicle-treated mice. The P161-PWY was among the top 5 pathways most strongly correlated with circulating IAA levels in ILA-treated mice (Fig. 4J, while very few pathways correlated with circulating ILA Fig. S2G). When including both ILA- and vehicle-treated mice, the P161-PWY pathway was the most strongly correlated with IAA levels (Fig. S2F). P161-PWY activity was also positively correlated with the cumulative EAE score and associated with a lower ILA:IAA ratio (Fig. 4K-L).

### IAA supplementation inhibits remyelination and is toxic to differentiating OLs in vitro

To investigate the functional relevance of changes in IAA levels on remyelination, we supplemented CPZ mice with 2mM IAA or vehicle in a similar experimental setup as for ILA earlier (Fig. 5A). Two weeks into the remyelination (and treatment) phase there was less remyelination of the CC with IAA supplementation compared to vehicle (Fig. 5B). The number of mature OLs was reduced with IAA, while the staining for the OPC marker PDGFRα was unchanged (Fig. 5C). *In vitro* IAA blocked OPC differentiation (Fig. 5E) and had toxic effects on differentiating OLs (Fig. 5F). IAA had no significant effect on markers of immune activation and reactive gliosis in the CC (Fig. 5D), which aligns with our observation that IAA supplementation had no effect in the neuroinflammation-driven EAE model. Here, IAA did not affect peak scores and, if at all (not significant with n=20 per group), trended towards worsened recovery (Fig. S3).

**Fig. 5.**
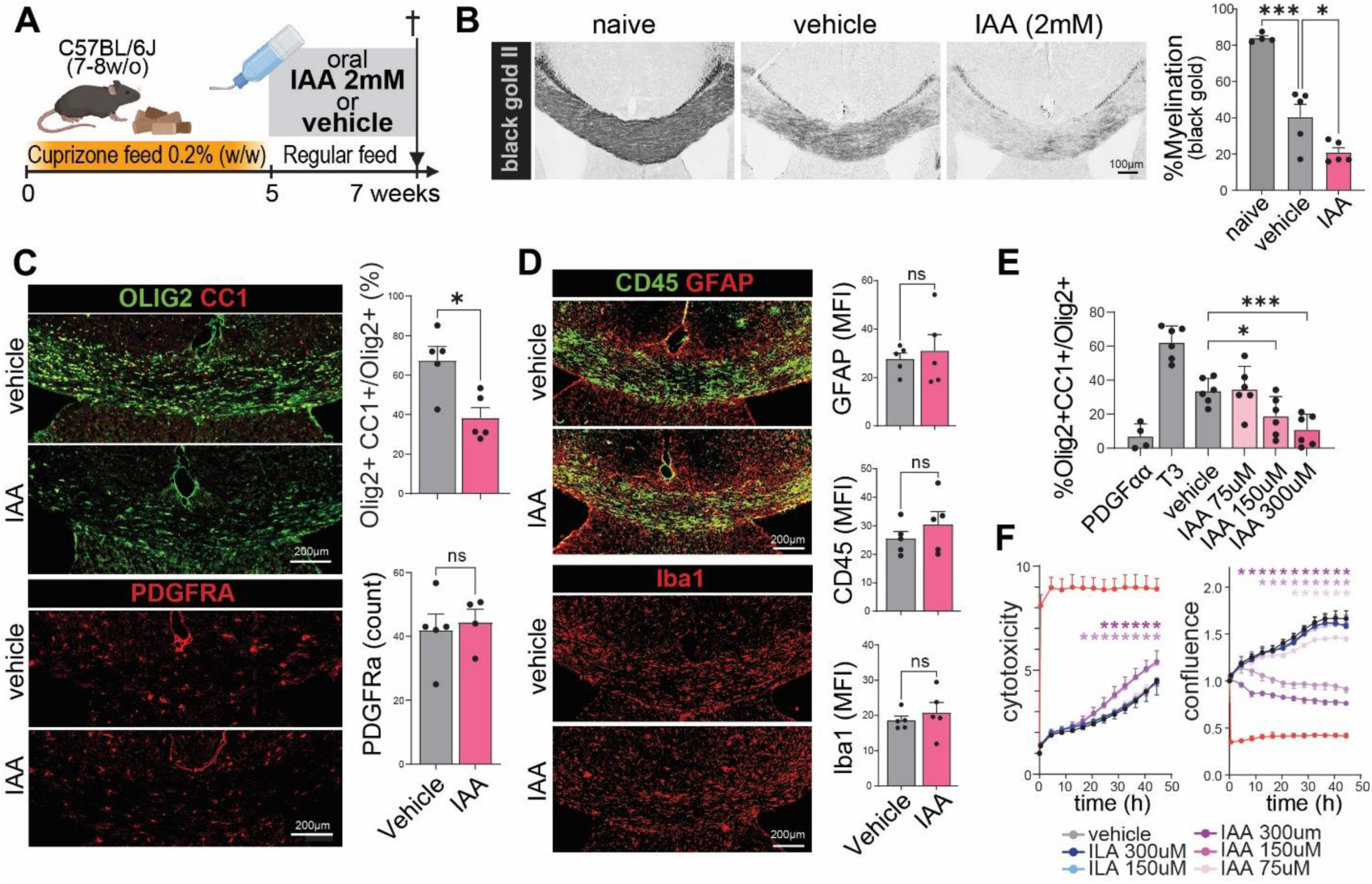
IAA supplementation inhibits remyelination and is toxic to differentiating OLs *in vitro*. (A) Schematic of the experimental setup, where IAA (2mM) or vehicle was supplemented during the remyelination phase. (B) Histopathology analysis of the corpus callosum (CC) myelination after two weeks of treatment with representative images (left) and quantification of the mean % myelination (right). (C-D) Representative images of immunofluorescence staining for mature oligodendrocytes (Olig2+CC1+) and OPCs (PDGFRα+), leukocytes (CD45), astrogliosis (GFAP), and monocytes/macrophages and microglia (Iba1) in the CC after two weeks of treatment with quantification of the % mature oligodendrocytes of all oligodendrocyte lineage cells (%Olig2+CC1+/Olig2+), the OPC count and the mean fluorescent intensities (MFI) for all remaining markers. n=5 mice per group for B-D. (E) Quantification of the percentage of primary murine OPCs differentiated into mature oligodendrocytes (%Olig2+CC1+/Olig2+) in the presence of vehicle or 75-300uM IAA *in vitro* (same setup as in Fig. 3A) (n= 6 independent experiments). (F) *In vitro* live cell imaging assays of primary murine OPCs treated with IAA (shades of purple), ILA (shades of blue), vehicle (black), or lysis buffer (red; positive control for cell death). Shown is the cell permeability dye uptake into differentiating OPCs i.e. cytotoxicity (left) and the confluence (right) over 48h normalized to baseline measurements for each well (shown is one representative of three independent experiments). Asterisks indicate significance at the corresponding time point compared to the vehicle and the asterisks’ colors represent the metabolite and its concentration at which the significant effects are observed. All error bars in the figure represent ±SD from the mean. Statistical significance was determined by one-way ANOVA with Dunnett’s post-hoc test for B and E and by Mann–Whitney U test for C-D (*P <0.05; ***P <0.001). For F multiple t-tests were performed comparing each metabolite condition to vehicle and the Holm-Šídák method was used to adjust for multiple comparisons (P <0.05).

### Imbalance of ILA and IAA levels (lower ILA and higher IAA) is associated with MS severity

We and others have shown that ILA is lower in pwMS, while IAA is not significantly different (*3, 4*). In the current study, we investigated the role of these metabolites in MS disease severity and progression using several independent cohorts. In the adult International Multiple Sclerosis Microbiome Study (iMSMS) cohort (n=345 with scoring), higher circulating IAA and a lower ILA:IAA ratio were associated with worse age-related MS severity (gARMSS) scores. In a second independent smaller JHU cohort (n=266), there was a similar trend (p-value=0.062) for the ILA:IAA ratio (Fig. 6A; Supplementary Tables Sheet 6). However, in this more extensively characterized cohort, circulating ILA and IAA correlated with multiple other clinical and patient-reported outcomes, with the ILA:IAA ratio having the most significant associations with these disease severity measures (Fig. 6B; Supplementary Tables Sheet 7). In a large multi-center cohort of pediatric-onset MS (POMS) with longitudinal follow-up (n=381), higher ILA levels were associated with reduced subsequent disease activity (new T2 lesions or new Gd-enhancing lesions). IAA levels showed no significant association with imaging outcomes, but a trend was observed with an increased annualized relapse risk (Fig. 6C; Supplementary Tables Sheet 8).

**Fig. 6.**
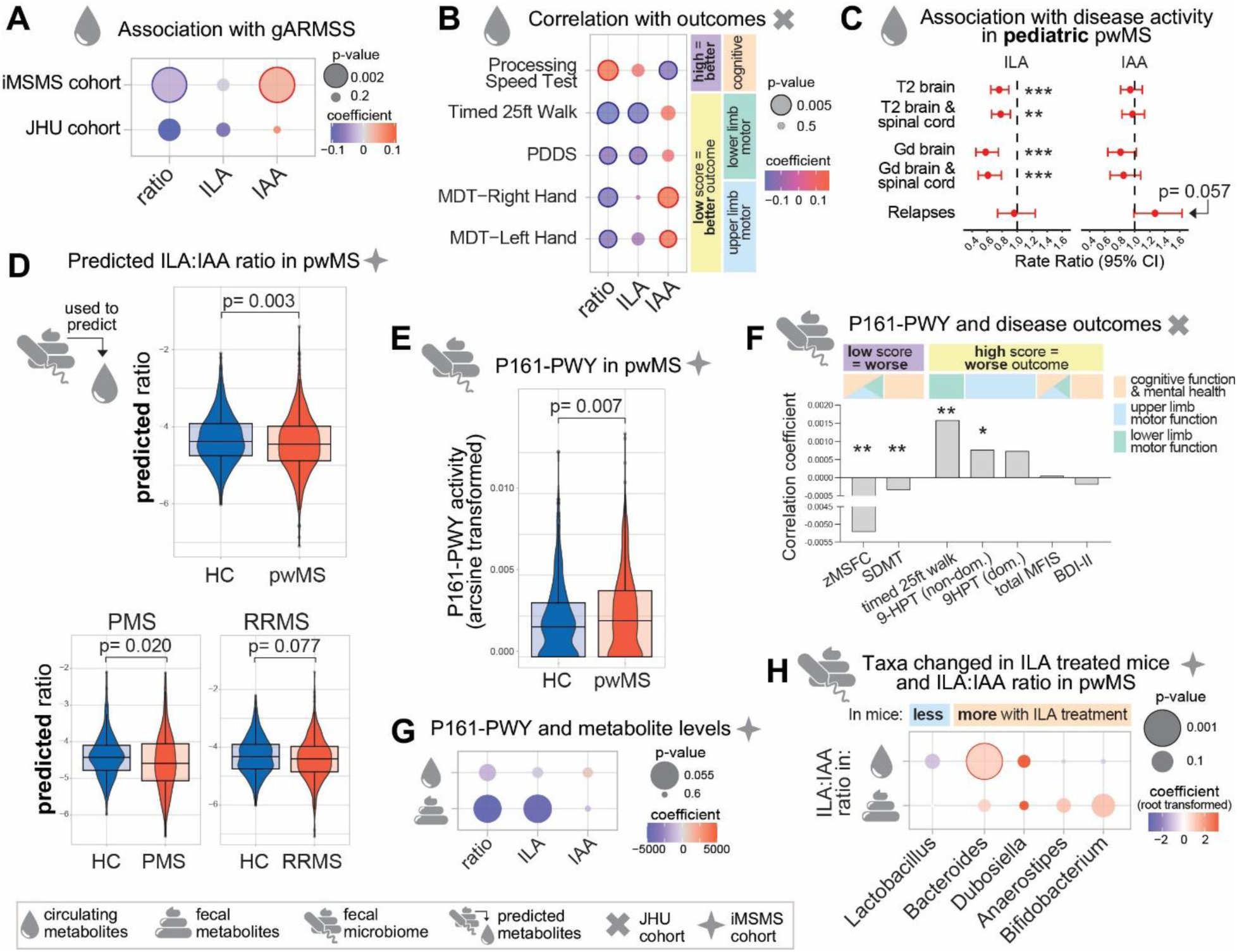
Gut microbiota functions altered by ILA supplementation are dysregulated in pwMS and the circulating ILA:IAA ratio is associated with MS outcomes and MS gut dysbiosis. (A) Association of circulating ILA, IAA and ILA:IAA ratio with the global Age-Related Multiple Sclerosis Severity Score (iMSMS cohort n=345 and JHU cohort n=266) with blue indicating negative and red positive association and dot size indicating the p-value (P <0.05 highlighted with solid border). (B) Correlation of the circulating metabolites with clinical and patient-reported outcomes collected for the JHU cohort (n=266) with dot color indicating the correlation coefficient and size indicating the p-value (P <0.05 highlighted with solid border). (C) Forrest plot showing incidence rate ratios of disease activity outcomes (annualized MRI lesions or clinical relapses) in POMS patients comparing high vs low ILA or IAA levels (median dichotomization) (n=381). (D) ILA:IAA ratios predicted based on the iMSMS(*15*) gut microbial taxonomy data in pwMS and household controls (n=576 households and progressive MS n=139 households and relapsing-remitting MS n=437 households). (E) Gut microbiota activity of P161-PWY, a functional pathway altered with ILA treatment, in pwMS and household controls from the iMSMS study(*15*) (n=576 households). Results are more significant when eliminating households with an inactive pathway (Fig. S4C) (F) Association of P161-PWY activity with clinical and patient-reported outcomes (n= 25-32; JHU cohort). (G) Association of P161-PWY activity with circulating and fecal metabolite levels in pwMS (n=696; iMSMS cohort). (H) Association of the genera changed by ILA treatment in mice with circulating and fecal metabolite levels in pwMS (n=696; iMSMS cohort) with blue indicating negative and red positive association (coefficients were root transformed for clearer visualization; untransformed results in supplementary table) and dot size indicating the p-value (P <0.05 highlighted with solid border). Statistical analyses were carried out with generalized linear models (for A, F-H), linear mixed-effect models (for D-E), and Spearman’s rank correlation tests (for B). All analyses were adjusted for age and sex and where available additionally BMI (all iMSMS cohorts and F) and race (all JHU cohorts). Multi-center data with household controls was additionally adjusted for household and study sites. (*P <0.05; **P <0.01; ***P <0.001).

### The predicted ILA:IAA ratio obtained from the microbiome composition is altered in pwMS

To investigate whether the changes in ILA and IAA in pwMS could be explained by changes in the gut microbiome, we calculated predicted ILA and IAA levels using the taxonomy data from the previously published iMSMS data (*15*) (n=576) using the online TrpNet tool (*16*). Predicted ILA levels were lower, and predicted IAA levels were higher in pwMS compared to healthy household controls (Fig. 6D, Fig. S4A), and those predictions correlated with experimentally determined values in both serum and stool (Fig. S4B).

### Gut microbiota functional pathways and taxa targeted by ILA supplementation are dysregulated in pwMS

Finally, we investigated if the taxa and microbial pathways changed with ILA treatment in mice are relevant targets in pwMS. The P161-PWY was up-regulated in pwMS compared to healthy household controls in the iMSMS dataset (Fig. 6E, the difference was even larger when only considering households where the pathway is active [activity>0] in both participants Fig. S4C). In a smaller JHU cohort subgroup (n=23-31) with metagenomics data, higher P161-PWY activity was associated with worse clinical and patient-reported outcomes (Fig. 6F; Supplementary Tables Sheet 9).

The P161-PWY activity positively correlated with circulating IAA levels in mice. In pwMS, there was a trend of an association of higher pathway activity with lower ILA levels and ILA:IAA ratio in the feces though this did not reach statistical significance (Fig. 6G; p-value for ILA =0.06; p-value for ratio =0.06; Supplementary Tables Sheet 10). When only including pwMS in whom this microbial pathway is active in the analysis, higher P161-PWY activity was significantly associated with a lower fecal ILA:IAA ratio (p-value=0.04; Supplementary Tables Sheet 10).

Several of the genera altered in ILA-treated mice have previously been shown to be altered in pwMS (*17–19*). Therefore, we focused on the associations of these genera with circulating ILA and IAA levels. Interestingly, a higher abundance of *Bacteroides* was associated with a higher circulating ILA:IAA ratio (and IAA) in pwMS (Fig. 6H). *Bifidobacterium* and *Lactobacillus* were associated with higher fecal ILA and IAA, respectively (Supplementary Tables Sheet 11). We observed no significant sex differences in P161-PWY activity, in circulating ILA, IAA levels and their ratio, and in the predicted metabolite levels.

## DISCUSSION

In pwMS, the circulating metabolome is dysregulated with an imbalance of gut-derived reductive and oxidative AAA metabolites. ILA (produced through a reductive pathway) is one of the top dysregulated metabolites. It is reduced in pwMS with particularly low levels in progressive MS.^1,2^ Here, we show ILA supplementation impacts biological processes relevant to MS, namely neuroinflammation and remyelination, and restores gut microbial oxidative/ reductive metabolic imbalance.

ILA reduced peak and cumulative EAE scores, which is most likely driven by its effects on peripheral immune cell activation and infiltration – both key processes in the model’s acute stage. Our *in vitro* studies suggest these could be direct effects of ILA on immune cells, particularly on T-cells by reducing their activation and polarization towards pathological Th17 cells and on myeloid cells by reducing their inflammatory activation. This is supported by previous reports of ILA’s anti-inflammatory effects on immune cells in the periphery, including T-cells (*8, 20, 21*), dendritic cells (*22*), monocytes and macrophages (*3, 21, 23, 24*). ILA’s effects on neuroinflammation alongside its effects on remyelination are particularly promising because neuroinflammation can jeopardize the success of other avenues of treatment, such as remyelination by blocking OPC differentiation.

A possible limitation of our *in vitro* results with immune cells and OPCs (discussed below) is the high ILA concentration required to achieve effects. In our study, we saw effects at 200-500uM ILA. This aligns with previous *in vitro* studies where at least 100uM up to 10mM ILA are required (*8, 21–23, 25*). Possible explanations for this could be that ILA is internalized or catabolized warranting either a constant supply of low ILA levels as it is physiologically the case *in vivo* or a single treatment at higher concentrations to ensure enough ILA is available for the *in vitro* experiment duration.

Another hallmark of MS besides neuroinflammation is demyelination and failure of adequate remyelination. ILA supplementation promoted remyelination in the CPZ-induced animal model of toxic demyelination. Our *in vitro* studies with both human and murine OPCs suggest this might be a direct effect of ILA on OPCs, promoting their differentiation. As OPC differentiation is impaired in pwMS (*26*), ILA’s capacity to promote OPC differentiation makes it a promising therapeutic candidate. However, it is unclear whether ILA’s remyelinating effects *in vivo* are exclusively mediated by promoting OPC differentiation. Besides OPC differentiation, numerous other factors impact remyelination, including the inflammatory milieu. In the CPZ model, ILA also reduced reactive gliosis and myeloid cell activation and increased homeostatic microglia staining. This suggests ILA may further indirectly impact remyelination by reducing the local inflammatory milieu and (based on our previous findings in human PBMCs (*3*)) by promoting phagocytosis of myelin debris. ILA’s dual effect on remyelination and inflammation could make it a very relevant remyelination agent that can overcome barriers of remyelination such as the CNS compartmentalized inflammation in MS.

Having established ILA’s effects on neuroinflammation and remyelination in animal models *in vivo* and on immune cells and oligodendroglia *in vitro*, we sought to determine its cellular target receptor. ILA’s effects on immune cells (and enterocytes) have previously been shown to be mediated by AhR (*23, 25*), epigenetic modifications (*22*), and in humans additionally by the HCA3 receptor (*21, 24*). Therefore, we focused on investigating ILA’s yet unstudied mechanism of action on oligodendrocyte lineage cells. Transcriptomic analysis showed enrichment of AhR-responsive pathways in OPCs with ILA treatment. AhR is a ligand-activated transcription factor that senses environmental and metabolic ligands and induces various ligand- and context-dependent responses ranging from toxicity to anti-inflammatory effects. It is expressed at low levels on murine and at higher levels on human OL lineage cells (*27*). Using AhR-/- cells and an AhR inhibitor, we demonstrated that ILA’s effects on both murine and human OPC differentiation *in vitro* were AhR-dependent. This is reassuring since the sensitivity to AhR-ligands varies across species (*28*). In the future, it would be interesting to further dissect the relevance AhR for ILA’s effects on OL cells *in vivo* using conditional oligodendrocyte-specific AhR-/- mice. For a such study, an inducible knock-out would be essential, as AhR-/- mice show myelination abnormalities during development (*29, 30*). However, experiments with an inducible oligodendrocyte-specific AhR-/- mouse, could be challenging as AhR deletion itself affects OL function and the susceptibility of OLs to inflammatory cytokines (*29, 31*). Furthermore, AhR is a pleiotropic receptor that initiates various downstream effects depending on the ligand (*32*). IAA may also act via this receptor (*33, 34*). Nonetheless, overall our findings and previous studies show ILA has direct, possibly redundant, effects on immune and glial cells, which are at least partially mediated by AhR.

We further investigated if ILA also impacts circulating IAA levels, because the change in circulating gut-derived metabolites in pwMS comes with an underlying shift of gut microbial AAA pathways (*3*). In both naive and EAE mice, ILA supplementation reduced circulating IAA levels, suggesting ILA restores the gut microbial oxidative and reductive indole metabolite imbalance.

Similar to ILA, circulating IAA is primarily produced by the gut microbiota (*6*). Therefore, we hypothesized that oral ILA supplementation might impact the gut microbiota and thereby alter circulating IAA levels. Indeed, ILA changed the gut microbiome’s alpha diversity, beta diversity, and functional pathways. The bacteria altered with ILA supplementation are associated with ILA and IAA production. In our mouse experiments, the abundance of *Bifidobacterium* and *Dubosiella* positively and of *Lactobacillus* negatively correlated with the circulating ILA:IAA ratio. In pwMS, higher *Bacteroides* abundance was associated with a higher circulating ILA:IAA ratio. Supporting this, *in vitro* studies have shown *B. thetaiotaomicron* produces larger quantities of ILA than IAA (*35*) and is one of the few species that can break down IAA to skatole (*36*). *Bifidobacterium* and *Anaerostipes* species have been shown to produce higher amounts of ILA than IAA or completely lack IAA-producing pathways (*35, 37*). Most of the targeted bacteria also have immunomodulatory effects. *B. thetaiotaomicron*, *D. newyorkensis* and *B. pseudolongum* restored the Treg/Th17 balance in a murine model of inflammatory bowel disease (*38–40*). In a rat EAE model *Bifidobacterium* supplementation reduced disease severity (*41*). Both *Dubosiella* and *Anaerostipes* have previously been shown to mildly negatively correlate with EAE severity (*42, 43*). Whether *Lactobacillus*, which was downregulated with ILA supplementation, is detrimental or beneficial in EAE or MS is unclear: *L. reuteri* increases CNS autoimmune susceptibility and EAE severity (*44, 45*), while *L. crispatus* and *L. rhamnosus* supplementation reduces EAE severity in rats (*41*) (though to a much lesser extent than *Bifidobacterium*). *Lactobacillus* produces ILA but also IAA, depending on the *Lactobacillus* species (*44, 46*), which unfortunately remained unidentified in our study. Interestingly gut taxa targeted by ILA supplementation are also dysregulated in pwMS. *Bacteroides* (in particular the *B. thetaiotaomicron* species) (*17, 18*), *Anaerostipes* (*18*), and *Bifidobacterium* (*17*) have been reported to be less abundant in pwMS while *Lactobacillus* is more abundant in pwMS (*19*).

The acetylene degradation pathway, P161-PWY, was the most significantly upregulated gut microbial pathway with ILA supplementation. One of the key enzymes of this pathway is aldehyde dehydrogenase (AldA). AldA has broad substrate specificity for different aldehyde substrates (*47*), including indole-3-acetaldehyde which it oxidizes to form IAA (*48*). This links the ILA-induced downregulation of the pathway with decreased circulating IAA levels. Indeed, the pathway activity was among the top three pathways most significantly correlated to circulating IAA levels in ILA-treated mice and in pwMS higher P161-PWY activity was associated with a lower fecal ILA:IAA ratio. In pwMS P161-PWY activity was higher than in healthy controls and was associated with worse outcomes. The most significantly upregulated pathways were tetrahydrofolate biosynthesis pathways (FOLSYN-PWY and PWY-6612), which are enriched in the young compared to the aged gut microbiome in mice (*49*) and promotes the survival of regulatory T-cells in the gut (*50*). This shows ILA supplementation alters the activity of MS-relevant gut microbial functional pathways.

Since ILA supplementation alters IAA-producing gut microbial taxa and pathways and reduces circulating IAA, we investigated the functional relevance of modulating IAA levels. In the CPZ model, IAA attenuated remyelination. This is most likely due to IAA’s detrimental effects on mature and maturing OLs, which we observed both *in vivo* (reduced mature OLs) and *in vitro* (toxic effects on differentiating OPCs). IAA had no significant effect on markers of reactive gliosis and immune cells, suggesting its effects in the CPZ model might primarily be on OLs. Supporting the notion that IAA has little effect on neuroinflammation, IAA had no effect on the clinical score in EAE. At first, these results may seem surprising in light of reports of IAA’s anti-inflammatory and anti-oxidative effects (*51–53*). However, IAA’s effects depend on the cell type and environment. In the absence of peroxidases, IAA exerts anti-inflammatory and anti-oxidative effects (*52, 54*), while in their presence IAA undergoes oxidative carboxylation to form free radicals (*55*). This causes oxidative stress and lipid peroxidation and explains IAA’s well-characterized cytotoxic effects, especially in cells with high peroxidase activity (*33, 56*). In MS and its preclinical models tissue peroxidase activity is increased through the inflammatory activation of local and infiltrating immune cells, contributing to increased free radical formation and oxidative stress (*57–59*). The increased vulnerability of OLs to oxidative stress and lipid peroxidation (especially in the CPZ model (*60*)), may explain IAA’s detrimental effect on these cells (*61, 62*).

Based on our findings that ILA supplementation impacts circulating IAA and IAA’s detrimental effects in preclinical models, we examined the relationship between IAA, ILA levels and ILA:IAA ratio with disease severity in pwMS. While ILA levels are reduced in pwMS, and associated with reduced disease severity (*3, 4*), IAA is not significantly altered and its association with disease severity is unclear. In this study IAA and ILA were associated with several distinct disease outcomes and the ILA:IAA ratio provided a good overall measure, significantly correlating with most outcomes. This highlights that the imbalance of ILA and IAA plays a role in MS disease progression. Since ILA supplementation both elevates circulating ILA levels and reduces circulating IAA levels, it is a promising approach to restoring this imbalance in pwMS. Using a microbiome-based metabolite prediction tool, we found that the gut microbiome in pwMS has an overall (predicted) reduced capacity to produce ILA while producing more IAA with their ratio being the most significantly altered. This shows that the MS-related gut dysbiosis impacts levels of both molecules. The rather weak correlation between taxa-based predicted levels and experimental levels could be due to a methodological limitation of the TrpNet tool or other factors, such as dietary uptake of metabolites, that also impacts fecal and circulating metabolite levels.

Interestingly, in POMS, IAA was not associated with disease activity, while higher ILA levels, in line with previous reports (*5*), were associated with reduced disease activity. One possible explanation may be deduced from our preclinical studies where ILA both alleviated neuroinflammation and promoted remyelination while IAA only impacted remyelination. Since POMS is driven more by inflammation (*63*) rather than failed remyelination (*64*), ILA with its anti-inflammatory effect might have a greater impact on disease activity in POMS. Besides this, the differences in POMS and adult MS, could also be explained by increasing IAA levels throughout life (*65*) (in our analyses, predicted IAA levels also increased with age in HCs ρ=0.143 [CI: 0.063-0.233] p-value=0.0006), which might make a disease-related further upregulation of IAA more relevant in older pwMS. This also suggests that ILA supplementation might be particularly beneficial in progressive MS and warrants further ILA supplementation studies in aged mice, as we might see an even greater treatment effect.

In summary, supplementation of ILA, a gut-derived metabolite that is reduced in pwMS, dampens neuroinflammation and promotes remyelination. ILA reduces circulating IAA, a remyelination-blocking metabolite associated with worse outcomes in pwMS, and alters gut microbial taxa and pathways regulating the ILA:IAA balance. These taxa and pathways are dysregulated in pwMS. Thereby this study not only identifies a novel remyelination agent but also provides mechanistic insights into how MS-associated gut dysbiosis may affect remyelination and MS disease progression.

## MATERIALS AND METHODS

Below a brief summary of the key methods. Please refer to the supplementary materials and methods for more details.

### Mice

C57BL/6J WT mice and 2D2 TCR mice were obtained from The Jackson Laboratory. Ahr-/- mice were generated by breeding Ahr^fx^ mice with CMV-Cre transgenic mice. Mice were kept in pathogen-free conditions with ad libitum food and water under a 12-hour light/dark cycle.

### Experimental autoimmune encephalomyelitis

EAE was induced in 10-12-week-old C57BL/6J mice via subcutaneous injection of 150 µg MOG35-55 peptide in CFA, with pertussis toxin administered on the day of immunization and 48 hours later. EAE progression was monitored daily using a clinical score from 0 (normal) to 5 (death). One week post-immunization, mice were randomized to receive either vehicle (acidified water) or ILA (2 mM) in drinking water. In parenteral ILA studies, ILA was administered at 10-20 mg/kg twice daily. Tissue was then harvested 4 weeks post-immunization for immunohistochemistry or 18 days post-immunization for flow cytometric analysis of the immune infiltrate at the peak of EAE. For flow cytometry, spinal cords were digested, dissociated, and centrifuged in Percoll. Cells were stained with surface and intracellular antibodies and analyzed on a Cytek Aurora flow cytometer. Data were analyzed in FlowJo.

### Cuprizone-induced Demyelination

7-8-week-old C57BL/6J mice were fed a 0.2% cuprizone diet for 5 weeks, followed by 2 weeks of normal chow to allow for remyelination. Mice were randomized into vehicle or ILA (or IAA; both at 2 mM in drinking water) groups during the remyelination phase.

### Tissue Processing, Staining, and Imaging

Mice were perfused with PBS followed by 4% PFA, and brain/spinal cord tissues were fixed overnight and cryoprotected in 30% sucrose. Tissues were embedded in O.C.T. and sectioned. Myelination was assessed using the Black Gold II (Biosensis) and FluoroMyelin (Thermo Fisher) staining kits on 20 µm sections, and quantification was performed using ImageJ. For immunofluorescence, sections were rehydrated, antigen-retrieved, and incubated with primary antibodies overnight, followed by secondary antibodies and Hoechst staining. Images were captured using a Zeiss Axio Observer Z1 and analyzed using ZEN and ImageJ software. For transmission electron microscopy (TEM), mice were perfused with glutaraldehyde-PFA fixative, and the corpus callosum was post-fixed in osmium tetroxide. Thin sections (60-90 nm) were prepared for TEM and analyzed for g-ratio and myelinated axons using MyelTracer software.

### In Vitro Experiments with T-cells and Macrophages

Macrophages were isolated from C57BL/6J mice injected with 1.5 mL of 3% sodium thioglycolate. After 3-5 days, peritoneal macrophages were harvested, plated, and allowed to adhere. Cells were stimulated with LPS (10 ng/mL) + IFN-γ (10 ng/mL) and treated with ILA or vehicle. The effect of ILA on NO production by activated peritoneal macrophages was measured using a Griess assay. Naive CD4+ T cells were isolated from the spleen of 2D2 TCR mice using a negative selection kit. For Th17 polarization assays the cell were then plated with whole recombinant MOG, irradiated splenocytes and a Th17 polarization cocktail. For proliferation and activation assays the cells were further stained with a proliferation dye after the isolation and then also plated with MOG and irradiated splenocytes. In both assays the wells were either treated with vehicle or various concentrations of ILA and cultured for 3 days. On day 4 the cells were stimulated with PMA/ionomycin, and proliferation as well as Th17 differentiation were analyzed by flow cytometry after staining for activation and Th17 markers.

### Murine and Human OPC Differentiation Assay

Murine primary oligodendrocyte precursor cells (OPCs) were isolated via immunopanning and cultured in proliferation media for 48 hours. The media was then replaced with control media containing vehicle or ILA (75, 150, 300 µM) for 48 hours. Cells were also treated with proliferation and differentiation media as controls. Differentiation was assessed by immunocytochemistry, RNA-Seq, qPCR, and live cell imaging. Human embryonic stem cells (hESCs) expressing a triple knock-in OPC reporter were maintained in mTeSR plus media. Differentiation was induced by neural induction medium supplemented with dual SMAD inhibitors and retinoic acid, followed by culture in N2B27 and PDGF media. At day 75, OPCs were purified by MACS using CD90.2 microbeads. For ILA treatment, purified OPCs (≥90% purity) were plated in PDGF media and treated with ILA. Nano-luciferase (Nluc) activity was measured at days 0 and 5 using a NanoGlo luciferase assay. MBP-Nluc activity was quantified as fold change in luminescence.

### Murine Gut Microbiome Analysis

C57BL/6J mice were immunized with MOG as described above to induce EAE and then treated with ILA or vehicle after 1 week. Fecal pellets were collected on days 0, 16, and 28, and plasma was collected on day 28. Fecal DNA was extracted, and 16S rRNA sequencing was performed using Illumina Miseq. Microbiome data were analyzed for diversity and differences between groups using R and microbiomeMarker. Functional analysis was performed using Picrust2 and Metacyc databases.

### Quantification of Indole Lactate and Acetate from Mouse Plasma

Plasma was collected from naïve mice after 24 and 72 hours of ILA supplementation, and from EAE-induced mice as part of the microbiome analysis. Plasma was stored at −80°C until processing. Metabolites were extracted using 80% methanol with a heavy isotope internal standard (serotonin-D4). After centrifugation, supernatants were dried and resuspended in water with formic acid, and analyzed by LC-MS/MS.

### Metabolomics and Metagenomics Analyses in Pediatric-onset and Adult MS

For the pediatric-onset multiple sclerosis (POMS) analyses, plasma was collected from individuals with MS onset before age 18, recruited from 17 U.S. sites (demographics in table). Metabolomics was carried out with untargeted GC-TOF MS. For adult MS analyses (demographics of the cohorts in table), blood and stool samples were collected and stored at −80°C. For the JHU cohort, plasma metabolomics was carried out using GC/MS or LC/MS/MS, and stool sample metagenomic sequencing was carried out by Innomics Inc. For the iMSMS cohort, both serum and fecal samples were analyzed by Metabolon Inc. The relationship between metabolite levels and clinical, patient-reported, or MRI outcomes was modeled using negative binomial regression, generalized linear models, and Spearman’s rank correlation.

## List of Supplementary Materials

***Supplementary Figures:*** Fig. S1-S4

***Supplementary Materials and Methods***

***Supplementary Tables:*** Sheet 1-11

*Sheet 1:* Differentially expressed genes in ILA-compared to vehicle-treated OPCs (bulk RNASeq)

*Sheet 2:* Differentially regulated pathways in ILA-compared to vehicle-treated OPCs (bulk RNASeq)

*Sheet 3:* Differentially abundant gut microbial genera with ILA treatment compared to vehicle (FDR <5%)

*Sheet 4:* Differentially abundant gut microbial functional pathways with ILA treatment compared to vehicle (FDR <0.5%)

*Sheet 5:* Demographics of patient cohorts

*Sheet 6:* Association of circulating metabolites with the global Age-Related Multiple Sclerosis Severity Score

*Sheet 7:* Association of circulating metabolites with patient-reported and clinical outcomes

*Sheet 8:* Effect of circulating ILA and IAA on the rate ratio of MRI disease activity and clinical relapse in POMS

*Sheet 9:* Association of the P161-PWY with patient-reported and clinical outcomes

*Sheet 10:* Association of the P161-PWY activity with fecal and circulating metabolite levels (in all pwMS) and Association of the P161-PWY activity with fecal and circulating metabolite levels (only in the subgroup of pwMS with P161-PWY activity)

*Sheet 11:* Association of taxa altered with ILA treatment in mice with fecal and circulating metabolite levels

## Acknowledgments

We gratefully acknowledge Dr. David Joel Hackam MD PhD for providing us with AhR-/- mice.

Further, we acknowledge the University of Iowa Holden Comprehensive Center’s Microbiome Core for their essential support in providing microbiome sequencing services.

## Funding

The National Multiple Sclerosis Society supported this work with a postdoctoral fellowship (FG-2207-40205) awarded to LJ, and a Harry Weaver Neuroscience Scholar Award (JF-2007-36755) to PB.

The NIH/NINDS supported PB (R21NS123141), EW (R01NS117541), XC (K00EY029011), SG (T32AI007260) and PAC (R01NS041435).

Additional support was provided by USDA Project 2032-51530-025-00D to JWN. The USDA is an equal-opportunity employer and provider.

## Author contributions

Conceptualization: PB, LJ, SS. Data curation: LJ, SG, FS, VAS. Formal analysis: LJ, SS, JL, AD, FS, SG, VAS, KS. Funding acquisition: PB, LJ, PAC, XC, SG, ELW, JWN. Investigation: LJ, SS, JL, AD, DJ, VM, KG, SG, PD. Methodology: LJ, SS, JL, AD, KG, SG, PD, FS, VAS, KS, PB. Project administration: PB, LJ, SS. Resources: PB, PAC, XC, MK, NH, AM, ELW, SEB, JN, KB. Software: LJ, FS, SG, KS, VAS. Supervision: PB, PAC, XC, MK, NM, AM, ELW, SEB, JN. Validation: LJ, SS, JL, AD, SG, PD. Visualization: LJ. Writing – original draft: LJ, SS. Writing – review & editing: PB and the final draft all authors.

## Competing interests

The authors declare that they have no competing interests in relation to this work, except for a pending patent (PCT Application No. PCT/US24/27239) concerning the application of ILA as a therapeutic agent for multiple sclerosis. The patent applicant is Johns Hopkins University, and the authors of the patent are Pavan Bhargava, Kathryn Fitzgerlad, Peter Calabresi and Michael Kornberg. This patent is currently under review.

## Data and materials availability

Additionally to the data are available in the main text and the supplementary materials, count/OTU tables and code for omics datasets generated for this study are deposited in the GitHub repository for this project (https://github.com/L-Jank/ILA-Supplementation-in-MS.git) which will be made public upon publication of this project.

## Supplementary Materials

### SUPPLEMENTARY FIGURES

**Figure S1:**
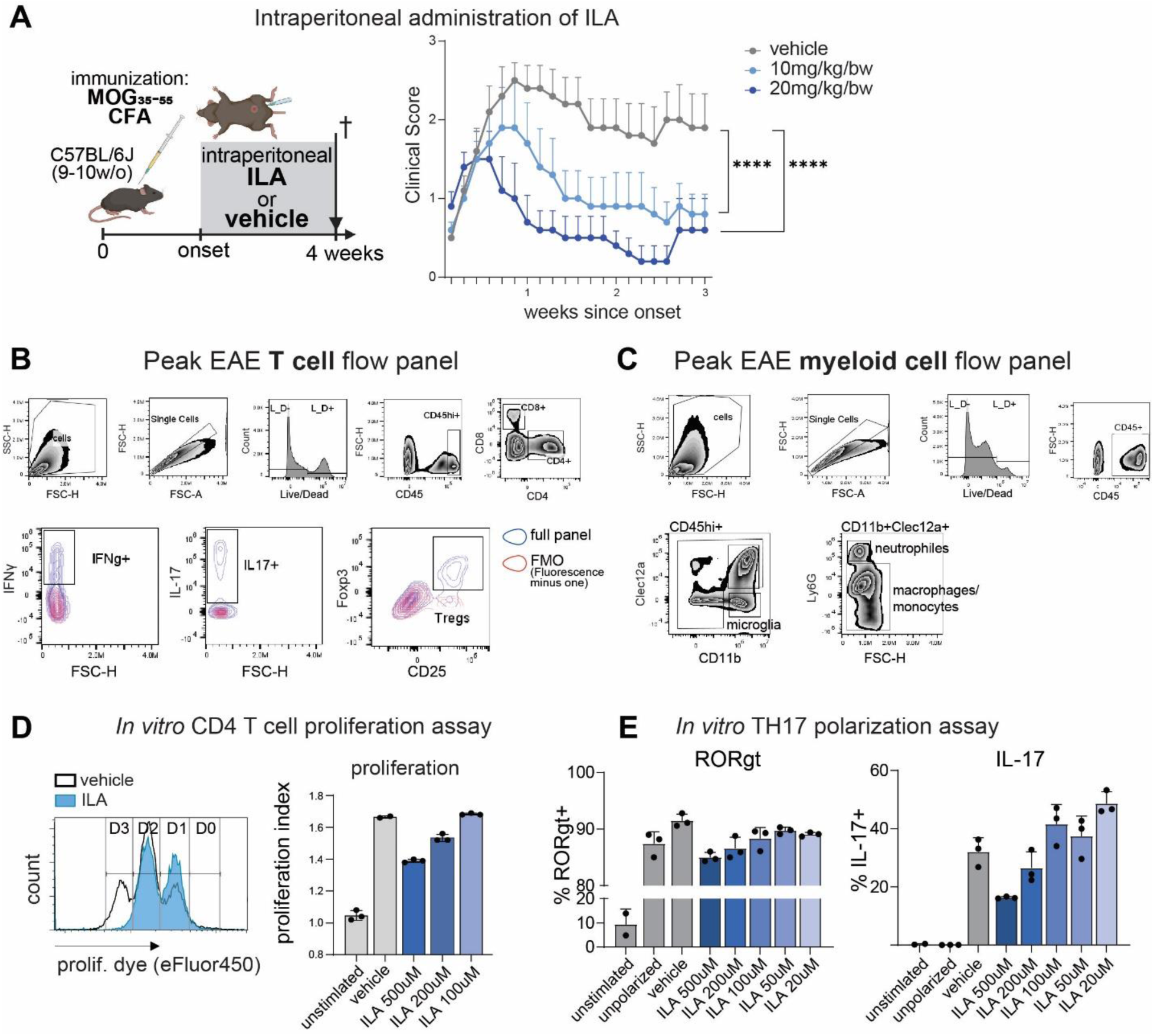
ILA supplementation attenuates EAE. (A) Schematic of the experimental setup (left) and Mean Hooke clinical EAE severity score of mice randomized at disease onset into 3 groups injected intraperitoneally twice daily with either vehicle or a low or high dose of ILA ±SEM (right). (B-C) Gating strategy for T cells and myeloid cells used in flowcytometric experiments determining ILA’s effects on the spinal cord immune infiltrate at peak of EAE (shown in Fig. 1E-G). (D-E) Flow cytometric analysis of T-cell proliferation and Th17 polarization assays with WT CD4+ T-cells. Shown are a representative histogram for the proliferation dye intensity in T cells treated with vehicle or ILA (D left), quantification of the mean proliferation index (D right), and the mean % of RORgt+ and IL-17+ T cells in the Th17 polarization assay (E) after three days of culture with vehicle or 20uM-500uM ILA. The n=5 mice per group for A and D-E show 3 technical replicates from one of two independent experiments. Error bars represent ±SD, unless mentioned otherwise. Statistical significance was determined by two-way ANOVA with Tukey’s multiple comparisons test for EAE scoring and one-way ANOVA with Tukey’s post-hoc test for D-E. (*P <0.05; **P <0.01; ***P <0.001; ****P <0.0001).

**Figure S2:**
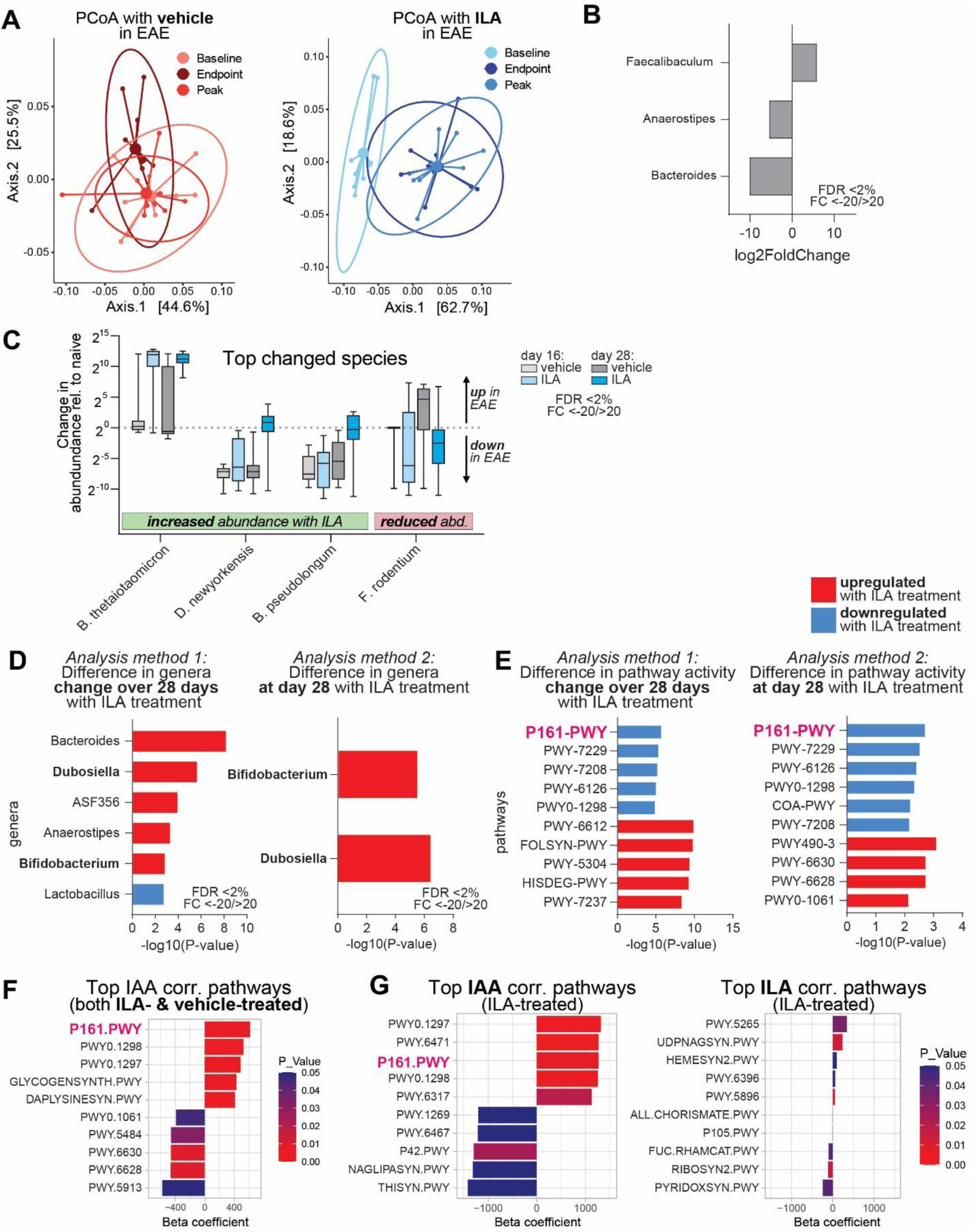
ILA supplementation alters the gut microbiome. (A) Principal coordinates analysis (PCoA) for both vehicle (left plot in red) and 1mM ILA (right plot in blue) with shades representing time points. (B) Gut microbial genera significantly different at baseline (cutoff: FDR <2% and Foldchange <-20/>20) (C) Top changed species with ILA treatment compared to vehicle ranked by magnitude of change with ILA (cutoff FDR<2%). Shown is the mean change in abundance relative to baseline (pre-immunization). (D) Analysis of significantly altered taxa with ILA treatment compared to vehicle with two alternative analysis approaches: Method 1 (analysis approach used for Fig. 4) comparison of the change in bacterial abundances from baseline to day 28 in vehicle and ILA treated mice; and method 2 direct comparison of bacterial abundances at day 28 between the two groups. The results of the two analyses overlap though less significantly changed genera were identified with method 2. Upregulated genera are shown in red and downregulated in blue (cutoffs: FDR <2% and Foldchange <-20/>20). (E) Comparing the same analysis approaches for the gut microbial functional pathway analysis. The P161-PWY is the most significantly down-regulated with both approaches. (F) Top 5 most significantly positively and negatively IAA correlated pathways in ILA- and vehicle-treated mice. P161-PWY is the most strongly positively IAA correlated pathway. (G) Top 5 most significantly positively and negatively IAA (left) and ILA (right) correlated pathways in ILA-treated mice side by side, showing that circulating ILA levels are less strongly associated with microbial pathways. The n= 9-10 mice per group. Statistical significance for B-C was determined using the R “limma” package where the p values were obtained from moderated t-statistic and adjusted using the Benjamini and Hochberg’s method. All correlations are Spearman’s rank correlations.

**Figure S3:**
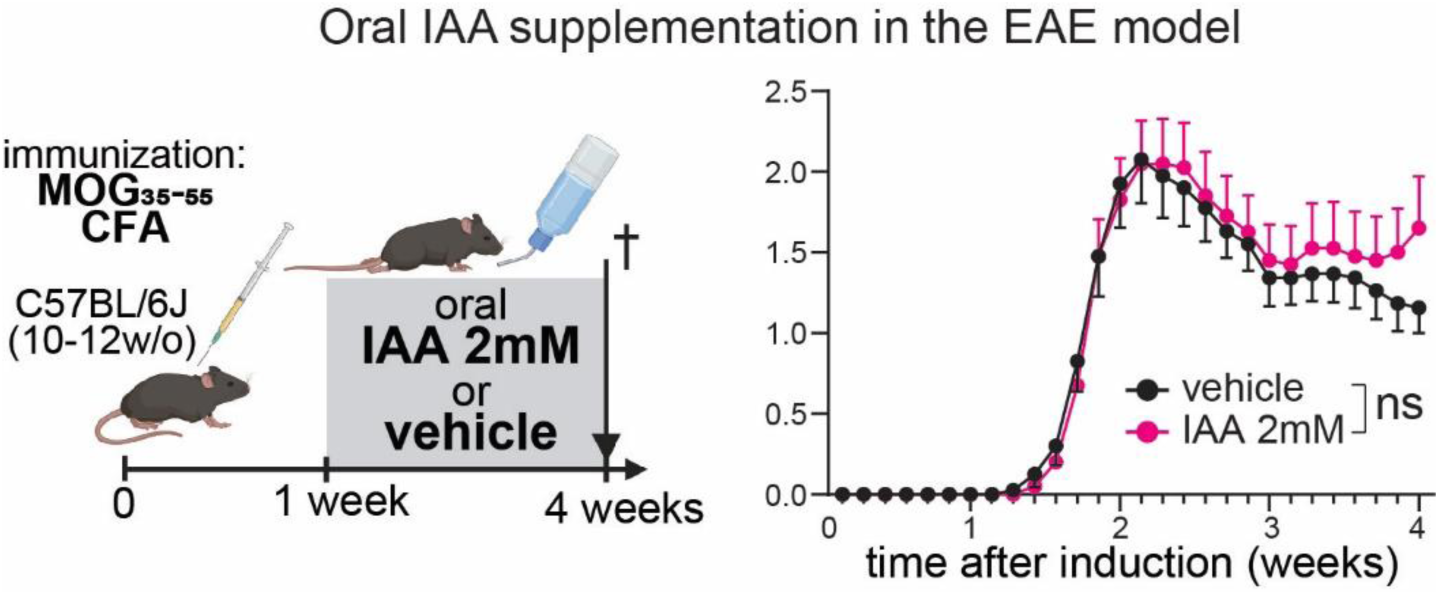
IAA supplementation has no significant effect on EAE severity. Schematic of the experimental setup (left) and Mean Hooke clinical EAE severity score of mice randomized 1 week post-immunization into a vehicle (acidified water) or ILA (2mM) group treated orally through the drinking water ±SEM (right). Statistical significance (here not significant) was determined by two-way ANOVA (n=20 mice per group from 2 experiments).

**Figure S4:**
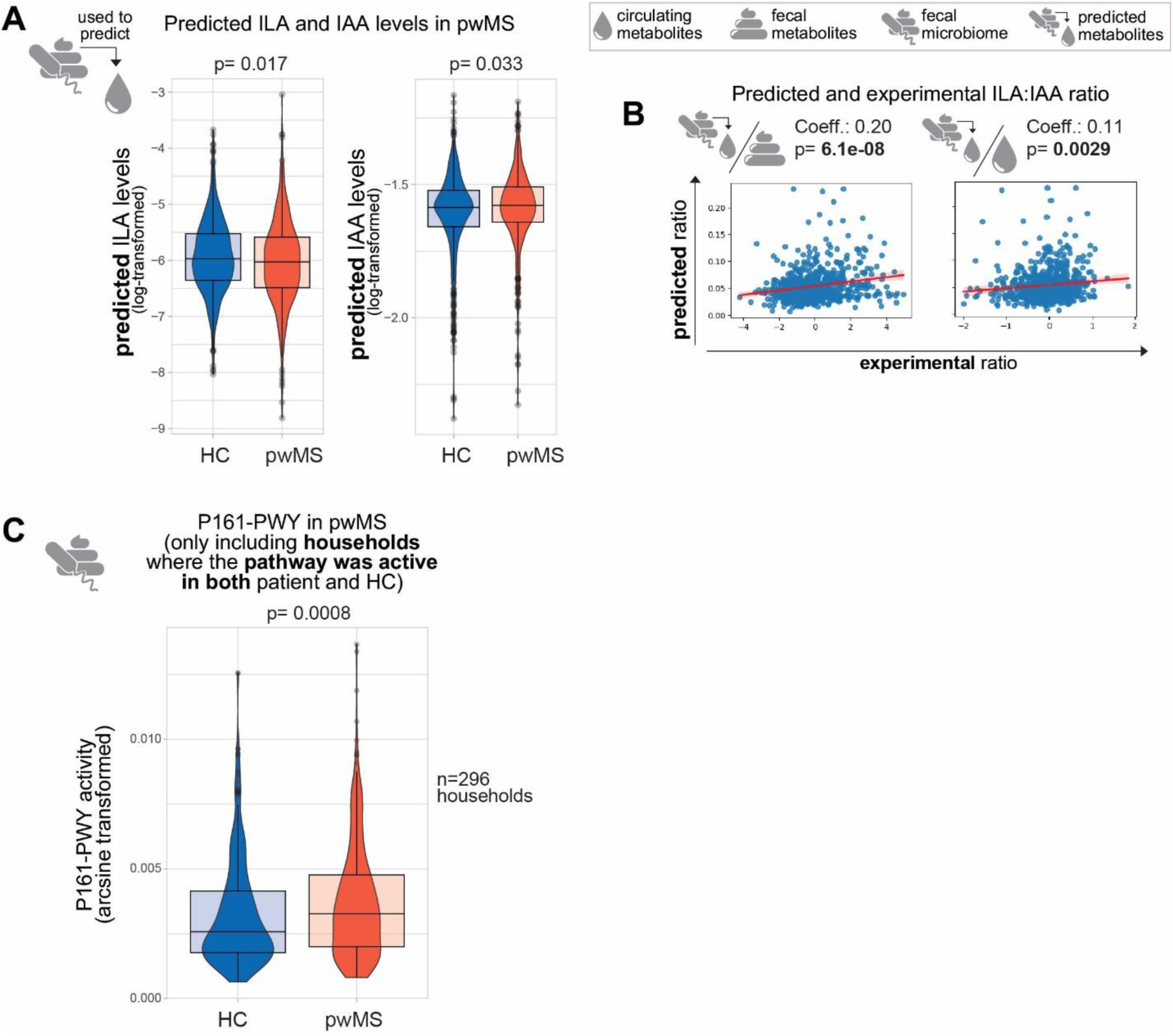
Gut microbiota functions altered by ILA supplementation are dysregulated in pwMS. (A) ILA and IAA levels predicted based on the iMSMS(*15*) gut microbial taxonomy data in pwMS and household controls (n=576 households). (B) Correlation of the predicted circulating ILA:IAA ratio to the experimental ratio in the feces (left) and serum (right) (n=576). (C) Gut microbiota P161-PWY activity in pwMS and household controls only including households from the iMSMS study (*15*) where the pathway was active in both the patient and healthy control (n=296 households). Statistical analyses were carried out with linear mixed-effect models for A and C, and Spearman’s rank correlation tests for B. All analyses were adjusted for age, sex and BMI. A and C were additionally adjusted for household and study sites.

### SUPPLEMENTARY MATERIAL AND METHODS

#### Mice

C57BL/6J mice and 2D2 TCR mice (C57BL/6-Tg(Tcra2D2,Tcrb2D2)1Kuch/J) were purchased from The Jackson Laboratory and housed in a pathogen-free animal facility at Johns Hopkins University School of Medicine. Global knockouts of *Ahr* (*Ahr^-/-^)* were kindly provided by Dr. David Joel Hackam MD PhD. As described previously(*66*), these mice were generated by breeding *Ahr^fx^* mice with transgenic mice expressing global Cre under the transcriptional control of a human cytomegalovirus minimal promoter CMV-Cre (B6.C-Tg(CMV-cre)1Cgn/J). All animals were kept under a 12-hour light/dark cycle, with ad libitum access to standard food and water, and housed in groups of 4-5 per cage. The use of experimental animals adhered to National Institutes of Health guidelines, and all methods and procedures were approved by the Johns Hopkins Institutional Animal Care and Use Committee.

#### EAE induction and ILA supplementation

Experimental autoimmune encephalomyelitis (EAE) was induced in 10–12-week-old male C57BL/6J mice through subcutaneous injection of 150 µg of MOG_35-55_ peptide (JHMI Peptide Synthesis Core, USA), emulsified in Complete Freund’s Adjuvant (CFA; Thermo Scientific, USA) containing 8 mg/ml of heat-killed *Mycobacterium tuberculosis* (BD, NJ). The injection was administered at two sites on the abdomen. To further enhance the immune response, 300 ng of pertussis toxin (List Biological Labs, California, USA) was administered intraperitoneally on the day of immunization, followed by a second dose 48 hours later.

The progression of EAE was monitored daily using a standardized clinical scoring system, ranging from 0 (normal) to 5 (death). Mice were scored as follows: a score of 0.5 indicated a limp tail tip, while a score of 1 indicated a fully limp tail. A score of 1.5 reflected limp tail and hind limb hindrance, and a score of 2 indicated hind limb weakness or dragging. Scores of 2.5 and 3 represented partial and complete hind limb paralysis, respectively, while a score of 3.5 indicated complete hind limb paralysis and forelimb weakness. A score of 4 signified complete hind limb and partial forelimb paralysis, and a score of 4.5 indicated complete paralysis of both the hind and forelimbs. A score of 5 was assigned in cases of death.

One week after immunization, the mice were randomly divided into two groups. The first group – the vehicle group – consisted of EAE mice that received acidified water (pH was adjusted to that of 2mM ILA). The second group – the ILA-treated group – received ILA (2mM, Sigma Aldrich) in their drinking water for the remainder of the experiment. The concentration was selected to deliver an approximate daily dose of 1.2mg per mouse, and water consumption was monitored by weighing the bottles daily.

In separate experiments involving parenteral ILA administration, mice were randomized at disease onset and received ILA injections twice daily, with doses adjusted to either 10mg/kg or 20mg/kg body weight.

At the end of the experiments, animals were perfused with PBS followed by PFA and brain and spinal cord tissues were collected for histological analysis, molecular studies, and flowcytometry as described in subsequent sections.

#### Cuprizone-induced demyelination and ILA supplementation

To induce demyelination, 7-8-week-old female C57BL/6J mice were fed a diet containing 0.2% Cuprizone (CPZ; Sigma-Aldrich, USA), a copper chelator, mixed into standard rodent chow for five consecutive weeks. During this period, all CPZ-fed animals were closely monitored for changes in weight, health status, and behavior, and fresh CPZ chow was provided three times per week. Control (naïve) mice were maintained on standard rodent chow for the duration of the experiment.

After five weeks of CPZ feeding, CPZ-fed animals were returned to a normal diet of standard chow for an additional two weeks to allow for the initiation of spontaneous remyelination. At the beginning of this remyelination phase, CPZ-fed animals were also randomized into two groups: 1) Vehicle groups: Animals on acidified water; 2) ILA treated group: animals treated with ILA (2 mM) added in drinking water for two weeks.

At the end of the experiments, animals were perfused and brains were collected for histological analysis as described in subsequent sections.

#### Brain and Spinal cord tissue processing

Mice were transcardially perfused with ice-cold 0.1 M phosphate-buffered saline (PBS), followed by 4% paraformaldehyde (PFA) in PBS. After perfusion, brain and spinal cord tissues were isolated and fixed in 4% PFA overnight at 4°C. The tissues were then transferred to 30% sucrose for cryoprotection over 72 hours. Once cryoprotected, the tissues were embedded in Optimal Cutting Temperature (O.C.T.) compound.

For histological analysis, brain sections were cut coronally at the level of the corpus callosum, and spinal cord sections were cut at a thickness of 20µm. For immunostaining, sections were prepared at 12µm thickness. All tissue sections were stored at −80°C until further processing.

#### Black Gold II Myelin Staining

The Black Gold II Myelin Staining Kit (Biosensis) was used to examine the demyelination. 20 µm coronal sections were warmed on the slide warmer for 30 min at 55-60°C, followed by staining with Black-Gold II, according to the manufacturer’s instructions. Stained brains and spine sections were imaged on Zeiss Axio Observer Z1 with multi-focal Z-point-supported tiling. Myelination was quantified by Image J software (https://imagej.nih.gov/ij/download.html). We conducted a quantitative analysis of the myelinated areas (corpus callosum for CPZ-model and spinal cord lesions and surrounding areas for EAE) by measuring the percentage of the field/area stained with Black-Gold II.

#### Immunofluorescence staining

Prior to immunostaining, cryostat sections of the spinal cord and brain were placed on a slide warmer for 30 minutes to dry. The sections were then rehydrated in phosphate-buffered saline (PBS) for 10 minutes. Antigen retrieval was performed using citrate buffer (pH 6.0), as needed depending on the specific antibodies.

Following antigen retrieval, sections were blocked with a blocking buffer containing 5% normal goat serum (NGS) in PBS and 0.4% Triton X-100 for 1 hour at room temperature. Primary antibodies, diluted in 3% NGS in PBS and 0.1% Triton X-100, were incubated overnight at 4°C. After several washes with PBS, sections were incubated with appropriate fluorophore-conjugated secondary antibodies for 1 hour at room temperature.

For nuclear staining, Hoechst 33342 (Thermo Scientific) was applied. The stained sections were then mounted using Aqua-Poly/Mount (Polysciences, USA). Imaging was conducted with a Zeiss Axio Observer Z1 epifluorescence microscope, and images were analyzed using ZEN 3.5 blue and ImageJ software. The antibodies used for immunofluorescence were: Iba-1 (FUJIFILM Wako, Poly Rabbit, #019-19741), GFAP (Agilent Dako, Poly Rabbit, #GA52461-2), Mac2 (Bio Legend, Mono Rat (M3/38), #125402), CD45 (Bio Legend, Mono Rat, #103112), TMEM119 (Synaptic System, Mono Mouse, #400 01), CD68 (Invitrogen, Mono Rat (FA-11), #14-0681-82), CC-1 (Millipore-Sigma, Mono Mouse, #OP80-100UG), Olig2 (Millipore-Sigma, Poly Rabbit, #AB9610), PDGFR-α (Cell Sig. Technology, Mono Rabbit, 3174S), MBP (EMD Millipore, Mono Rat, MAB386).

#### Transmission Electron Microscopy (TEM)

For TEM, mice were perfused with a fixative composed of 2.0% EM-grade glutaraldehyde, 2.0% EM-grade paraformaldehyde (PFA), and 3 mM magnesium chloride, combined with 0.05 M Sorenson’s phosphate buffer and 0.05 M sodium cacodylate buffer. Following perfusion, the brains were isolated and placed in freshly prepared fixative overnight. The corpus callosum was then carefully dissected and post-fixed in 2% osmium tetroxide (OsO₄) in a mixture of 0.075 M Sorenson’s phosphate buffer and 0.075 M sodium cacodylate for 2 hours at 4°C in the dark.

After rinsing the samples in distilled water they were dehydrated and embedded in EPON resin, followed by polymerization at 60°C overnight. Ultra-thin sections (60-90 nm) were prepared using a diamond knife on a Leica Ultracut UCT ultramicrotome. The grids were examined using a Hitachi 7600 transmission electron microscope (TEM) at 80 kV, and images were captured with an AMT CCD XR80 camera (8-megapixel, side mount, high-resolution, high-speed). Quantification of the g-ratio and myelinated versus unmyelinated axons was performed using MyelTracer(*67*) software

#### Flow cytometric analysis of the immune infiltrate at peak of EAE

Two independent flow cytometry experiments were conducted, each with 5 mice per group. In both experiments, 5 C57BL/6J mice per group were immunized with MOG35-55 peptide and sacrificed 18 days post-immunization. Non-immunized C57BL/6J mice served as controls. Mice were anesthetized with isoflurane, then perfused with cold HBSS. Spinal cords were dissected, chopped, and digested in a 37°C solution containing papain (20 U/ml) and DNase (100 U/ml) in papain buffer (22.5 mM D-glucose, 0.5 mM EDTA, 2.2 g/L NaHCO3, 5.5 mM L-cysteine in EBSS, pH 7.4). After 10 minutes of digestion with gentle rotation, tissues were mechanically dissociated with a 5 ml serological pipette. This process was repeated three times using progressively smaller pipette tips. Cells were filtered through a 70 µm filter, washed with PBS, and resuspended in 30% Percoll. After centrifugation at 550×g for 10 minutes, myelin debris was removed. Cells were washed in HBSS, incubated with Fc block (BioLegend 156604) and live/dead ZombieNIR (Thermo Fisher) in PBS for 15 minutes, then stained with cell surface antibodies (diluted 1:400) in FACS buffer for 30 minutes. For intracellular staining, cells were fixed with IC fix (eBioscience 88-8824-00) for 30 minutes, then stained with intracellular antibodies (diluted 1:200) in permeabilization buffer (eBioscience 88-8824-00) for 30 minutes. Cells were washed in FACS buffer and analyzed on a Cytek Aurora 3-laser (VBR) flow cytometer. Compensation was performed using single-stained UltraComp eBeads, fluorescent minus one (FMO) controls, and unstained or viability stain-only spinal cord cells. FlowJo software was used for analysis and gating (Fig. S1B-C). Antibodies used for the myeloid panel were: CD45-BV605, CD11b-PacificBlue, Clec12a-APC, I-Ab-PE, NOS2-PerCP-eF710, Arg1-PECy7, CD11c-BV570, Ly6C-FITC, Ly6G-BV785 and Zombie-NIR Live/Dead stain. The antibodies used for the lymphoid panel were: CD45-BV605, CD3-PerCP-Cy5.5, CD4-PE-Cy7, CD8-eF450, CD11b-BV510, IFNg-FITC, IL17a-APC, FoxP3-PE, CD25-eF610 and Live/Dead stain.

#### 2D2 TCR T-cell proliferation and Th17 polarization assays

8-12-week-old 2D2 TCR mice were euthanized, and spleens were harvested and placed into 5 mL of FACS buffer (PBS containing 2% FBS and 2 mM EDTA) on ice. The spleens were then mashed through a 100 µm wet filter using a sterile pestle to generate a single-cell suspension. After washing the filter with 10 mL of FACS buffer, the resulting cell suspension was collected into a 50 mL conical tube and centrifuged at 350g for 5 minutes to pellet the cells. The cell pellet was resuspended in RBC lysis buffer and incubated for 6 minutes on ice to lyse red blood cells. After lysis, the cell suspension was filtered through a 70 µm mesh and washed with FACS buffer. The final cell suspension was resuspended in 0.5 mL of FACS buffer per spleen, and the cell concentration was determined. Naive CD4+ T cells were isolated using the negative selection Naive CD4+ T Cell Isolation Kit (StemCell Technologies, #19765) according to the manufacturer’s protocol. The resulting enriched naive CD4+ T cells were centrifuged at 400g for 5 minutes and resuspended in FACS buffer. The purity of the naive CD4+ T cells was assessed by flow cytometry using the following antibodies: CD4-PE (clone RM4-5, Invitrogen), CD8a-PECy7 (clone 53-6.2, BioLegend), CD3e-FITC (clone 145-2C11, BD), CD11b-BV510 (clone M1/70, BioLegend), CD19-PacificBlue (clone 6D5, BioLegend), NK1.1-APC (clone PK136, BD), and propidium iodide. For proliferation and activation assays, the cells were washed twice with warm PBS and resuspended at 20 × 10^6 cells/mL in 0.5 mL PBS. The cells were incubated with eFluor 450 proliferation dye for 10 minutes at 37°C in the dark. After incubation, 5 volumes of cold CRPMI (RPMI-1640 with 10% FBS, 1% L-glutamine, 0.01 mM 2-mercaptoethanol, and 1% penicillin/streptomycin) were added, and the cells were incubated on ice for 5 minutes. The cells were then centrifuged at 500g for 5 minutes, washed three times with 5 mL of cold CRPMI, and resuspended in CRPMI at the desired concentration for downstream assays. For the proliferation and activation assays, 1 × 10^6 cells/mL of these labeled cells were plated with recombinant whole MOG (20 μg/mL) and irradiated splenocytes in CRPMI for 3 days. For Th17 polarization assays, 1 × 10^6 cells/mL of unlabeled naive CD4+ T cells were incubated with recombinant whole MOG (20 μg/mL), TGF-β (10 ng/mL), IL-1β (10 ng/mL), IL-23 (10 ng/mL), IL-6 (10 ng/mL), anti-IL4 (10 µg/mL), and anti-IFNγ (10 µg/mL) and irradiated splenocytes in CRPMI for 3 days. In both assays wells were additionally either incubated with vehicle or various concentrations of ILA. On day 4, the cells were resuspended in fresh media and stimulated with PMA and ionomycin (with Brefeldin and Monensin) for 5 hours. The cells were then stained for activation and Th17 markers and analyzed on a Cytek Aurora flow cytometer. The data was analyzed using FlowJo. Proliferation was assessed by measuring the proportion of proliferating cells.

#### Measuring NO production by activated peritoneal macrophage

10–12-week old C57BL/6J mice were intraperitoneally injected with 1.5 ml of 3% sodium thioglycolate. After 3-5 days, peritoneal macrophages were harvested from the peritoneal cavity with chilled Dulbecco’s phosphate-buffered saline (DPBS). The cells were centrifuged at 400g for 7 min and resuspended in the DMEM/F12-10 medium. We allowed the cells to adhere for 2-3 hours after plating and washed off any contaminated cells with PBS. Subsequently, the cells were stimulated with the vehicle or LPS (10ng/mL) +IFN-γ (10ng/mL) in the presence of various concentrations of ILA or vehicle. Then a Griess assay (Griess assay kit, Invitrogen) was performed with the supernatants according to the manufacturer’s instructions.

#### OPC differentiation assays with primary murine OPCs

Primary OPCs were isolated using immunopanning as previously described(*68*). In brief, the day before isolation, three 15 cm uncoated petri dishes were pre-incubated overnight at 4°C in Tris-HCl pH 9.5: two with 6 μg/mL goat anti-rat IgG, and one with 2.5 μg/mL BSL1. On isolation day, secondary antibody plates were washed and incubated with 1 μg/mL rat anti-mouse CD11b or PDGFRα in 0.2% BSA for 2 hours at room temperature. Neonatal forebrains from C57BL/6J mouse pups (P6-P9) were dissected and dissociated using 2.11 mg/mL papain and 100 U/mL DNase I in papain buffer. Cells were resuspended in 0.2% BSA, filtered, and subjected to a series of panning steps: BSL1 plate (10 min), CD11b plate (20 min), and PDGFRα plate (90 min). Bound cells were harvested using 0.0625% trypsin and plated on poly-d-lysine-coated surfaces. Cells were maintained in OPC basal media containing. Dulbecco’s Modified Eagle Medium high glucose (Thermo Fisher 31053028) containing 100U/100μg/ml penicillin/streptomycin (Fisher Scientific 15-140-122), 5μg/ml N-acetyl-l-cysteine, 1xSATO (1μg/ml human apo-transferrin, 1μg/ml BSA, 0.16μg/ml putrescine, 0.6mg/ml progesterone, 0.4ng/ml sodium selenite), 1xB27 (ThermoFisher 17504044), 1xTrace Elements B (Cellgro 99-175-C), 5μg/ml insulin, 10ng/ml d-biotin, 4.2μg/ml forskolin, 1mM sodium pyruvate (Millipore Sigma S8638), 4mM l-glutamine (Thermo Fisher 25-030-081). The proliferation media further contained the following growth factors: 40ng/ml PDGFAA (PeproTech 100-13A), 1ng/ml NT-3 (PeproTech 450-03), 10ng/ml CNTF (PeproTech 450-13).

For the differentiation assays the isolated OPCs were kept in OPC proliferation media for 48 hrs. Then the media was then completely replaced to control media containing 8 ng/ml PDGFaa and 10ng/ml CNTF, with vehicle or ILA (75, 150 300 uM) for an additional 48 hrs. Non-differentiating control media was supplemented with 40 ng/ml PDGFaa, 1ng/ml NT3 and 10ng/ml CNTF and differentiating control media was supplemented with10ng/ml CNTF and 40ng/ml T3. Readouts for these differentiation assays included immunocytochemistry, bulk RNASeq, qPCR and live cell imaging.

#### Immunocytochemistry of murine OPCs

For immunocytochemistry, the OPCs were cultured on poly-D-lysine pre-coated coverslips at a density of 15,000 cells per coverslip. After the assay, the coverslips were fixed in 4% paraformaldehyde (Sigma Aldrich P6148), permeabilized with 0.5% Triton, and blocked in 5% donkey serum. They were incubated overnight at 4°C with primary antibodies (rabbit Olig2 (Millipore AB9610), mouse CC1 (Calbiochem OP80), rat MBP (Millipore MAB386)), washed and then incubated with secondary antibodies (donkey anti-rabbit AlexaFluor 488 (Jackson ImmunoResearch 711-545-152), donkey anti-mouse AlexaFluor 555 (Invitrogen A-31570), donkey anti-rat AlexaFluor 647 (Jackson ImmunoResearch 712-605-153)) for 1 hour. After washing, cells were stained with DAPI (Invitrogen 21306) and mounted with Prolong Glass Antifade Mountant (Thermo Scientific P36984).

#### qPCR and bulk RNASeq analysis of primary murine OPCs

For RNASeq and qPCR the OPCs were plated in poly-d-lysine pre-coated 12 well plates at a density of 75.000 cells per well. At the end of the assay, the cells were lysed and RNA was isolated using the manufacturer’s instruction of RNeasy Micro Plus kit. RNA quality and quantity were assessed using a NanoDrop spectrophotometer. For the qPCR analysis 500ng of RNA was then used to make cDNA using iSCRIPT cDNA Synthesis Kit (Bio-Rad). The qPCR was performed using iQ SYBR Green reagent (Bio-Rad) on the CFX384 Touch Real-Time PCR Detection System (Bio-Rad). The delta-delta Ct method was used to normalize to the data (*β-actin* was used as a reference gene). The sequences for the primers used are: *β-actin* forward ACCTTCTACAATGAGCTGCG and reverse CTGGATGGCTACGTACATGG; *Mbp* forward TACCTGGCCACAGCAAGTAC and reverse GTCACAATGTTCTTGAAG; Cyp1a1 forward CATCACAGACAGCCTCATTGAGC and reverse CTCCACGAGATAGCAGTTGTGAC; Cyp1b1 forward GCCACTATTACGGACATCTTCGG and reverse ACAACCTGGTCCAACTCAGCCT.

Bulk RNASeq was carried out by Novogene. In brief, following RNA isolation, library preparation, and sequencing, transcripts were quantified using Salmon (*69*) and then imported into R with tximport (*70*). The data was then deconvoluted to correct for batch effects between experiments with ComBat (*71*) and differential genes were identified with DESeq2 (*72*). Up and downregulated pathways were identified using the EnrichR (*73*) and the WikiPathways (*74*) gene sets.

#### Live cell imaging of primary murine OPCs

Live cell imaging was carried out as described previously(*68*). In brief, primary murine OPCs were plated in 96-well plates at a density of 30,000 cells/well and cultured in proliferation media for 1-2 days. The cells were then treated with ILA, IAA or vehicle as indicated. Cytotox Green permeability dye (1:800; Sartorius 4632) was added to monitor viability. The cells were then placed in a Sartorius Incucyte live cell imager and imaged every 4 h. At the conclusion of the experiments, Incucyte software was used to measure the percent confluency and Cytotox green uptake, these values were then normalized to baseline measurements for each wells.

#### Oligodendrocyte differentiation from human embryonic stem cells and purification of the human reporter OPC

Human Embryonic Stem cell (hESC) line H9 (WA09; WiCell) was maintained in mTeSR plus media (Stemcell Technologies) in normal culture condition of 37 °C and 5% CO2. These cells were genetically-engineered to express triple knock-In OPC reporter system: PDGFR-TdTom-Thy1.2, MBP-secNluc and PLP1-sfGFP reporter.

hESC colonies were dissociated using Accutase and plated on Matrigel-coated plates at 120,000 cells per well of a 6-well plate in mTeSR Plus media supplemented with 5 mM blebbistatin. Once the colonies reached the desired size (approximately 2 days post-plating), differentiation was initiated by adding neural induction medium supplemented with dual SMAD inhibitors (SB431542 at 10 µM, LDN193189 at 250 nM), and all-trans RA at 100 nM. From day 8 to day 12, the media for differentiating cells were supplemented with RA (100 nM) and SAG (1 µM). On day 12, adherent cells were detached mechanically and cultured in low-attachment plates to allow cell aggregates to form spheres. The differentiating cells were maintained in N2B27 medium until day 20, after which PDGF media was used. On day 30, spheres were plated into poly-L-ornithine/laminin-coated 6-well plates in PDGF media. Two-thirds of the old media was carefully replaced with fresh PDGF medium every 2-3 days until day 75 of differentiation.

On day 75, differentiating cells were dissociated using Accumax. The single-cell suspension was passed through a ∼70 µm cell strainer (BD Biosciences), washed, and resuspended in MACS buffer (Miltenyi Biotec) for MACS-based cell sorting. MACS purification was performed following the manufacturer’s instructions using CD90.2 (THY1.2) microbeads (Miltenyi Biotec). Briefly, the single-cell suspension was incubated with microbeads at room temperature for 15 minutes to allow cell binding. The cell suspension was then passed through the magnetic column where the tdTomato+ OPCs remain bound. Following three washes with MACS buffer reporter OPCs were eluted with MACS buffer by applying pressure with the provided syringe.

#### Testing the effect of ILA on human reporter OPCs

MACS-purified reporter OPCs with a purity of ≥ 90% were used for the assay. Roughly 4K cells purified OPCs were plated in PLO-laminin-coated 384-well plates with 50 µL of PDGF media per well. Drug treatment was performed 2 days after plating. Prior to drug treatment, 40 µL of media was removed from each well for baseline measurement of Nluc activity (day 0 reading). Drug dispensing was carried out using an ECHO 555 acoustic liquid handler. The cells were then cultured in a 37°C, 5% CO2 incubator. After 5 days, 40 µL of media was collected from each well for day 5 Nluc reading. MBP-Nluc activity was measured using NanoGlo luciferase assay reagents (Promega N1150). A 2.5 µL aliquot of the NanoGlo reaction mix (substrate: buffer = 1:50) was diluted with an equal volume of water (1:1 dilution). Subsequently, 5 µL of the diluted NanoGlo reaction mix was added to each well containing 20 µL of media collected from day 0 and day 5, respectively, in separate white-bottom 384-well plates. Relative luminescence units (RLU) were measured with a microplate reader (Clariostar, BMG Labtech) at 460 nm with 0.1-second exposure. Reported MBP-Nluc activity was quantified as the fold change between day 5 and day 0.

#### Murine gut microbiome analysis

10–12-weeks aged C57BL/6J mice (10 mice per group) were immunized with MOG35-55 peptide and then randomized into two ILA-treated and two vehicle-treated cages after 1 week. Clinical EAE scoring and weights were monitored daily. Fecal pellets were collected before EAE induction (day 0), at the peak of EAE (day 16), and at the endpoint (day 28). For fecal pellet collection, mice were held by hand and gently pressed in the lower abdomen area. then, pellets were collected in a sterile tube and stored at −80°C for downstream processing. On day 28 we also collected plasma to quantify circulating ILA and IAA levels in these mice (details below).

A total of 57 samples (samples from three time points for 10 vehicle- and 9 ILA-treated mice) were analyzed for the gut microbiota abundance comparison. Fecal DNA was isolated using (please mention the kit you used) and 16S rRNA was sequenced using Illumina Miseq by amplifying V3-V4 region and the resulting *fastq* sequences were processed to generate amplicon sequence variants (ASVs) table using *dada2*(*75*) and naïve Bayesian classifier with the Silva 138 database in the Microbiome Core at Holden Comprehensive Cancer Center at the University of Iowa. The ASV table was further analyzed in R v4.2.0 using *phyloseq*(*76*). Samples had a total of 2730183 reads with a median of 46393. These samples were analyzed for alpha diversity, and then, those taxa whose reads sum in at least 10% of the sample was less than 100 were filtered out followed by normalization using median sequence depth. Beta diversity was calculated using Bray-Curtis dissimilarity metric and plotted as a PCoA plot. Microbiota differential analysis was performed at genus level using the *microbiomeMarker*(*77*) and *limma*(*78*) packages in R. The differential analysis to identify taxa altered with ILA treatment was carried out using two alternative analysis approaches: Method 1 (analysis approach used for Fig. 4) comparison of the change in bacterial abundances from baseline to day 28 in vehicle and ILA-treated mice; and method 2 direct comparison of bacterial abundances at day 28 between the two groups (Fig. S2D). The results of the two analyses overlapped though less significantly changed genera were identified with method 2. The cutoffs selected for this analysis were an FDR<2% and a foldchange of <-20/>20. For functional analysis using the *picrust2* (*79*) and Metacyc (*80*) database the same 57 samples as above were analyzed. Samples had a total of 21289500 reads with a median of 3706830.91. Those functional categories whose reads sum in at least 10% of the sample were less than 100 were filtered out followed by normalization using median sequence depth. The same two analysis approaches as described above for the gut microbiota abundance comparison were used to compare ILA- and vehicle-treated mice (Fig. S2E) with the P161-PWY being consistently among the top downregulated pathways with ILA treatment.

#### Extraction and quantification of indole lactate and acetate from mouse plasma by liquid chromatography-tandem mass spectrometry (LC-MS/MS)

For Liquid chromatography-tandem mass spectrometry (LC-MS/MS), we collected plasma (terminally by cardiac puncture) in naïve mice supplemented with 2mM ILA for 24 and 72hrs to study the change in circulating metabolites over time with ILA supplementation. In a second experimental setup we collected plasma from the EAE-induced mice included in the microbiome analysis described in more detail above. After collection, the plasma was stored at −80°C for downstream processing. To measure the ILA and IAA levels in the plasma, the metabolites were extracted using 1 mL of 80% ice-cold methanol (0.5% 1N HCl) containing pre-spiked heavy isotope internal standard serotonin-D4 (50 ng/mL). Samples were vigorously mixed for 5 min using a TissueLyser LT (Qiagen, Switzerland) at 50Hz, and centrifuged at 13000 rpm at 4 °C for 20 min. The supernatants containing metabolites and internal standards were collected and evaporated to complete dryness using a vacuum concentrator (Thermo Scientific “Savant SpeedVac” SPD 120P2, Waltham, MA, USA). The residues were then resuspended in 150 µL of 50% deionized water (ddH2O) containing 0.1% formic acid. Chromatographic separations of metabolites from the extracts achieved on a Kinetex Pentafluorophenyl (PFP/F5) stationary phase column (Phenomenex, Torrance, CA, USA) using a binary gradient mobile phase program (acetonitrile-eluent A and ddH2O-eluent B; both contained 0.1% FA) with the following gradient program: 0 – 2.1 min 100 % eluent B, 2.1 – 8 min, eluent A increased from 0% to 100%, 8 – 15 min eluent A hold at 100%, 15 – 15.1 eluent B increased to 100% from 0%, and hold for 3 min to allow for column equilibration before the next sample is injected using a Shimadzu ultrafast liquid chromatography (Shimadzu, Kyoto, Japan). The column was operated at 40°C with a constant mobile phase flow rate at 200 μL/min. Eluted compounds were introduced into Quadrupole Ion trap mass spectrometer (API4000 QTRAP LC-MS/MS, AB Sciex, ON, Canada), equipped with electrospray ionization (ESI) negative mode where individual metabolites were detected as [M-H]-molecular ions. An eight-point calibration curve of indole lactate standard in neat solvent is constructed to ensure instrument performance and linearity. Instrument control and quantification were performed using Analyst 1.7.1 and MultiQuant software (AB Sciex, Thornhill, ON, Canada) respectively.

#### POMS cohort metabolomics methods and analysis

Individuals with MS onset before the age of 18 years from seventeen sites in the U.S. Network of Pediatric were included if they were within four years of symptom onset, met the 2010 McDonald criteria for MS, were MOG IgG negative, and were followed for at least one year. Each collaborating site obtained human subject research approval through their respective ethics review committees, and all participants/ their legal guardians provided written informed consent allowing the use of their medical records for research.

Blood samples of University of California San Francisco (UCSF) participants were processed within 3 hours of collection. Samples from other 16 sites were drawn at the participating site and shipped overnight to UCSF, where samples were centrifuged and plasma was stored at −80°C. Frozen plasma was then aliquoted and shipped overnight on dry ice to the University of California Davis West Coast Metabolomics Center Central Services Core. Untargeted metabolomic profiling was performed in all samples as a batch to provide semiquantitative analyses of primary metabolism. Metabolites of primary metabolism were measured using untargeted GC-TOF MS based method (*81*) commercially available at the West Coast Metabolomics Center located at UC Davis. In short, column: Restek corporation Rtx-5Sil MS (30 m length x 0.25 mm internal diameter with 0.25 μm film made of 95% dimethyl/5%diphenylpolysiloxane); Mobile phase: Helium; Column temperature: 50-330°C Flow-rate: 1 mL min-1; Injection volume: 0.5 μL; Injection: 25 splitless time into a multi-baffled glass liner; Injection temperature: 50°C ramped to 250°C by 12°C s-1; Oven temperature program: 50°C for 1 min, then ramped at 20°C min-1 to 330°C, held constant for 5 min. Mass spectrometry parameters were used as follows: a Leco Pegasus IV mass spectrometer was used with unit mass resolution at 17 spectra s-1 from 80-500 Da at −70 eV ionization energy and 1800 V detector voltage with a 230°C transfer line and a 250°C ion source. Data processing: Raw data files were preprocessed directly after data acquisition and stored as ChromaTOF-specific *.peg files, as generic *.txt result files and additionally as generic ANDI MS *.cdf files. ChromaTOF vs. 2.32 was used for data preprocessing without smoothing, 3 s peak width, baseline subtraction just above the noise level, and automatic mass spectral deconvolution and peak detection at signal/noise levels of 5:1 throughout the chromatogram. Apex masses were reported for use in the BinBase algorithm. Result *.txt files were exported to a data server with absolute spectra intensities and further processed by a filtering algorithm implemented in the metabolomics BinBase database. The BinBase algorithm (rtx5) used the settings: validity of chromatogram (<10 peaks with intensity >10^7 counts s-1), unbiased retention index marker detection (MS similarity>800, validity of intensity range for high m/z marker ions), retention index calculation by 5th order polynomial regression. Spectra were cut to 5% base peak abundance and matched to database entries from most to least abundant spectra using the following matching filters: retention index window ±2,000 units (equivalent to about ±2 s retention time), validation of unique ions and apex masses (unique ion must be included in apexing masses and present at >3% of base peak abundance), mass spectrum similarity must fit criteria dependent on peak purity and signal/noise ratios and a final isomer filter. Failed spectra were automatically entered as new database entries if s/n >25, purity <1.0 and presenced in the biological study design class was >80%. All thresholds reflect settings for ChromaTOF v. 2.32. Quantification was reported as peak height using the unique ion as default, unless a different quantification ion was manually set in the BinBase administration software BinView. A quantification report table was produced for all database entries that are positively detected in more than 10% of the samples of a study design class (as defined in the miniX database) for unidentified metabolites. A subsequent post-processing module was employed to automatically replace missing values from the *.cdf files. Replaced values were labeled as ‘low confidence’ by color coding, and for each metabolite, the number of high-confidence peak detections was recorded as well as the ratio of the average height of replaced values to high-confidence peak detections. These ratios and numbers were used for manual curation of automatic report data sets to data sets released for submission.

The association between clinical (relapse count) and MRI (count of MRIs with at least one new T2 lesion or Gd+ lesion) outcomes with each metabolite (high vs. low levels, median dichotomization) was estimated using negative binomial regression models with a follow-up time offset (i.e., annualized rates). Adjustment for sex, age, self-reported race (White/other), biological mother’s highest degree, DMT usage (naive/lower efficacy/higher efficacy), BMI (underweight or healthy weight/overweight/obese/severely obese), site (UCSF vs non-UCSF), and number of available MRIs (for imaging-related outcomes) was performed.

#### JHU adult MS cohort metabolomics and metagenomics methods and analysis

A total of 266 participants were recruited from several recently completed clinical research studies at Johns Hopkins University (JHU). Human subject research approval was obtained through their respective ethics review committees, and all participants/ their legal guardians provided written informed consent allowing the use of their medical records for research. Blood samples were collected for metabolomics analysis and participants underwent comprehensive clinical assessments to evaluate disease severity during their study visits. These assessments included the Expanded Disability Status Scale (EDSS) (available for a subset of participants, n=196), processing speed tests, and the Multiple Sclerosis Functional Composite (MSFC; n=23), which comprises the Timed 25-Foot Walk test (T25-FW; n=27), the 9-Hole Peg Test (9HPT; n=25) administered either on a board or tablet, and the Symbol Digit Modalities Test (SDMT; n=31). Additional information included the Patient Determined Disease Steps (PDDS), the Modified Fatigue Impact Scale (MFIS; n=30), and the Beck Depression Inventory-II (BDI-II; n=30). The number of participants for all outcomes are also listed in Supplementary Tables Sheet 5.

For metabolomic analysis blood samples were processed within three hours of collection following a standardized protocol used across all studies at Johns Hopkins University. Serum or plasma samples were aliquoted and stored at −80°C until further analysis. At the end of each study, plasma samples were analyzed for metabolites by Metabolon Inc. (Durham, North Carolina) using their standardized methods using either gas chromatography coupled with mass spectrometry (GC/MS) or liquid chromatography coupled with tandem mass spectrometry (LC/MS/MS). Mass spectra obtained from these analyses were compared to a reference library for compound identification. The relative abundance of metabolites was quantified by calculating the area under the curve of the corresponding mass spectra.

For metagenomics participants from the JHU cohort were provided with OMNIgene GUT OMR-200 stool collection kits (DNA Genotek Inc., Ottawa, Ontario, Canada) and instructed to collect stool samples at home following standard procedures. Collected samples were stored at −80°C until shipped on dry ice to Innomics Inc. (Cambridge, MA, USA) for further processing. Microbial DNA was extracted from stool samples, and short-read libraries were prepared for metagenomic sequencing using a standardized in-house protocol at Innomics Inc. Genomic DNA was first fragmented to appropriate sizes, followed by size selection to ensure fragments of optimal length for sequencing. Fragment ends were repaired, and adenosine residues were added to the 3’ ends to facilitate ligation of sequencing adapters. After adapter ligation, a second round of size selection was conducted to further refine fragment sizes. Libraries were then amplified through PCR, and quality control (QC) was performed to assess library size distribution and concentration prior to sequencing. The prepared libraries were sequenced on the DNBSEQ-G400 platform (Innomics Inc., Cambridge, MA, USA), generating paired-end reads of high quality. Sequencing data were processed to remove low-quality and contaminant reads using SOAPnuke (BGI, China), with the following filtering criteria: reads with ≥50% adapter sequence similarity, allowing up to three mismatches, were discarded. Reads shorter than 150 base pairs were removed, as were those with ≥0.1% ambiguous bases (N content). Furthermore, reads in which ≥50% of bases had a Phred quality score below 20 were excluded. Clean reads, representing high-quality sequences, were retained for downstream analyses. The output quality scores were reported using the Phred+33 scale. In silico separation of bacterial reads from contaminant reads was performed using *Kneaddata* (*82*) to discard low-quality sequences. Sequencing reads were mapped by using *Bowtie2* (*83*) to remove contaminations. To characterize the microbial composition at the species level, taxonomic profiling was conducted using *MetaPhlAn*4 (*84*). This tool aligns metagenomic reads to a curated set of clade-specific marker genes. Additionally, a secondary taxonomic analysis was performed using *Kraken2* (*85*), employing the *PlusPF* database (available from: https://benlangmead.github.io/aws-indexes/k2). Functional profiling of the microbial communities was carried out with *HUMAnN*3 (*82*), which facilitated the identification and quantification of microbial gene families and metabolic pathways, providing insights into the functional potential of the microbiome.

#### iMSMS/ UCSF adult MS cohort with metabolomics and microbiome data description, methods and analysis

The microbiome data set for the iMSMS cohort shown in this study has previously been published.(*15*) In this manuscript, we further show novel metabolomics data (fecal and circulating ILA and IAA levels) from this iMSMS cohort. For both datasets, human subject research approval was obtained through their respective ethics review committees, and all participants/ their legal guardians provided written informed consent allowing the use of their medical records for research.

For both the microbiome and metabolomics analysis blood and stool samples were collected. Participants received stool sample collection kits and were instructed to collect two consecutive stool samples (with three vials: a dry Q-tip, a snap-frozen vial, and a vial with Luria-Bertani broth and 30% glycerol) at home. Samples were frozen for at least 12 hours before shipping with ice packs, and upon receipt, they were stored at −80 until further processing. Blood samples were collected during the initial visit. The blood was centrifuged at 2200g for 20 min and the serum layers were collected and stored at −80°C.

For metabolic profiling, fecal (150g/sample) and serum (150ul/sample) samples were analyzed by Metabolon Inc. (Durham, North Carolina) using their standardized methods. Metabolon analyzed and quality-controlled the raw data using proprietary software.

The metabolomics data was then further analyzed using a Generalized Linear Model (GLM) represented by the equation: 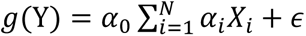 where Y is the response variable (analyte concentration) following a Gamma distribution, g(•) is the log link function, α₀ is the intercept, αᵢ are the model parameters, Xᵢ represent the predictor variables (severity score, age, BMI, gender, and site), N is the number of predictor variables, and ε is the residual term. This model allows for the analysis of the relationship between the analyte concentration and various clinical and demographic factors while accounting for the non-normal distribution of the metabolomic data.

## Notes

### Competing Interest Statement

The authors declare no conflicts of interest in relation to this work, except for a pending patent (PCT Application No. PCT/US24/27239) concerning the application of ILA as a therapeutic agent for multiple sclerosis. The patent applicant is Johns Hopkins University, and the authors of the patent are PB, KCF, PAC and MK. This patent is currently under review.

### Summary of Updates

Revised shortened version of the manuscript.

